# Differential impact of cell wall antibiotics on the Rod complex and aPBPs in *Bacillus subtilis:* Insights into the peptidoglycan elongation machineries

**DOI:** 10.64898/2026.07.24.740396

**Authors:** Charlène Cornilleau, Claire-Jing Rouchet, Aurélien Barbotin, Louise Destouches, Armand Lablaine, Elda Bauda, Cécile Morlot, Cyrille Billaudeau, Rut Carballido-López

**Affiliations:** Université Paris-Saclay, INRAE, AgroParisTech, Micalis Institute, 78350, Jouy-en-Josas, France; Univ. Grenoble Alpes, CNRS, CEA, IBS, Grenoble, France

**Keywords:** peptidoglycan, antibiotics, cell wall elongation machineries

## Abstract

Bacterial cell wall (CW), primarily composed of the biopolymer peptidoglycan, serve as essential protective barriers against external stresses and the internal turgor pressure. The peptidoglycan (PG) biosynthetic pathway encompasses sequential enzymatic reactions in the cytoplasm and in the membrane that involve critical enzymes susceptible to antibiotic targeting. Virtually each step of the pathway is the target of a known antibiotic. Antibiotic-induced inhibition of PG assembly typically weakens the sacculus, often leading to cell lysis. However, the cascade of events that follow inhibition of a specific enzyme of the pathway, and how these culminate in cell death remain largely unknown. Here, we investigated the effects on growing *Bacillus subtilis* cells of two categories of CW antibiotics: inhibitors of the synthesis of soluble PG precursors in the cytoplasm (fosfomycin and D-cycloserine) and inhibitors of the polymerisation and crosslinking reactions at the outer leaflet of the membrane, which incorporate newly externalised precursors into the existing network (vancomycin and penicillin). In *B. subtilis,* the latter reactions are catalysed along the sidewalls by the Rod complex, thought to primarily build the sacculus, and by class A penicillin-binding proteins (aPBPs), thought to add to repair it. Our findings reveal that the two antibiotic groups lead to growth arrest, sacculus thinning, and eventual cell lysis. However, while the impact of vancomycin and penicillin G is rapid, lacking morphological deformation, fosfomycin and D-cycloserine induce cell widening and bulging before lysis. During shortage of PG precursors, dysregulated PG hydrolytic activity contributes to elevated cell lysis but is not responsible of bulging. Instead, dispersed PG synthesis by aPBPs persists while the activity of the Rod system is rapidly arrested, resulting in cell rounding. We propose that this facilitates the redirection of the limited PG precursors to sites of CW repair, thereby preserving cell integrity and allowing for prolonged growth during antibiotic challenge.

## Introduction

Most bacteria are encased in a cell wall (CW) that protects them from external injuries and counteracts their very high internal osmotic pressure. The major structural component of the CW is the polymer peptidoglycan (PG), a load-bearing meshwork of glycan strands formed by repetition of GlcNAc-MurNAc (N-acetyl-glucosamine—N-acetyl-muramic acid) disaccharides that are cross-linked together by stem peptides containing L- and D-amino acids (Vollmer, Blanot, and De Pedro 2008; Pazos and Peters 2019). Such polymeric mesh, named the sacculus, gives rigidity and structural integrity while providing plasticity to allow expansion and division during growth. Because PG is essential and common to the majority of bacteria but is absent in higher organisms, the PG biosynthetic pathway is the most prominent target of antibiotics.

The PG biosynthetic pathway involves a concerted set of sequential enzymatic reactions that take place first in the cytoplasm (synthesis of soluble PG precursors), then in the inner leaflet of the membrane (synthesis of lipid-linked intermediates lipid I and lipid II) and finally in the outer leaflet of the membrane (polymerisation into the existing sacculus) (**Fig. 1**). Virtually each step is catalysed by an essential enzyme that is targeted by a known antibiotic (Sarkar et al. 2017). Lipid II is flipped to the outer leaflet of the cytoplasmic membrane by dedicated flippases and glycan strands are then polymerised by enzymes possessing transglycosylase (TG) activity while peptide bridges are formed through transpeptidation (TP) reactions (Sauvage et al. 2008). TG reactions are mediated by two types of polymerase systems: bifunctional penicillin-binding proteins of Class A (aPBPs), which possess both TG and TP activity, and SEDS (shape, elongation, division, sporulation)-family proteins. The TG activity of SEDS proteins is associated to the TP activity of Class B penicillin-binding proteins (bPBPs) within multi-protein complexes that consist of several integral membrane proteins and are associated to cytoskeletal polymers (**Fig. 1**).

**Fig. 1.**
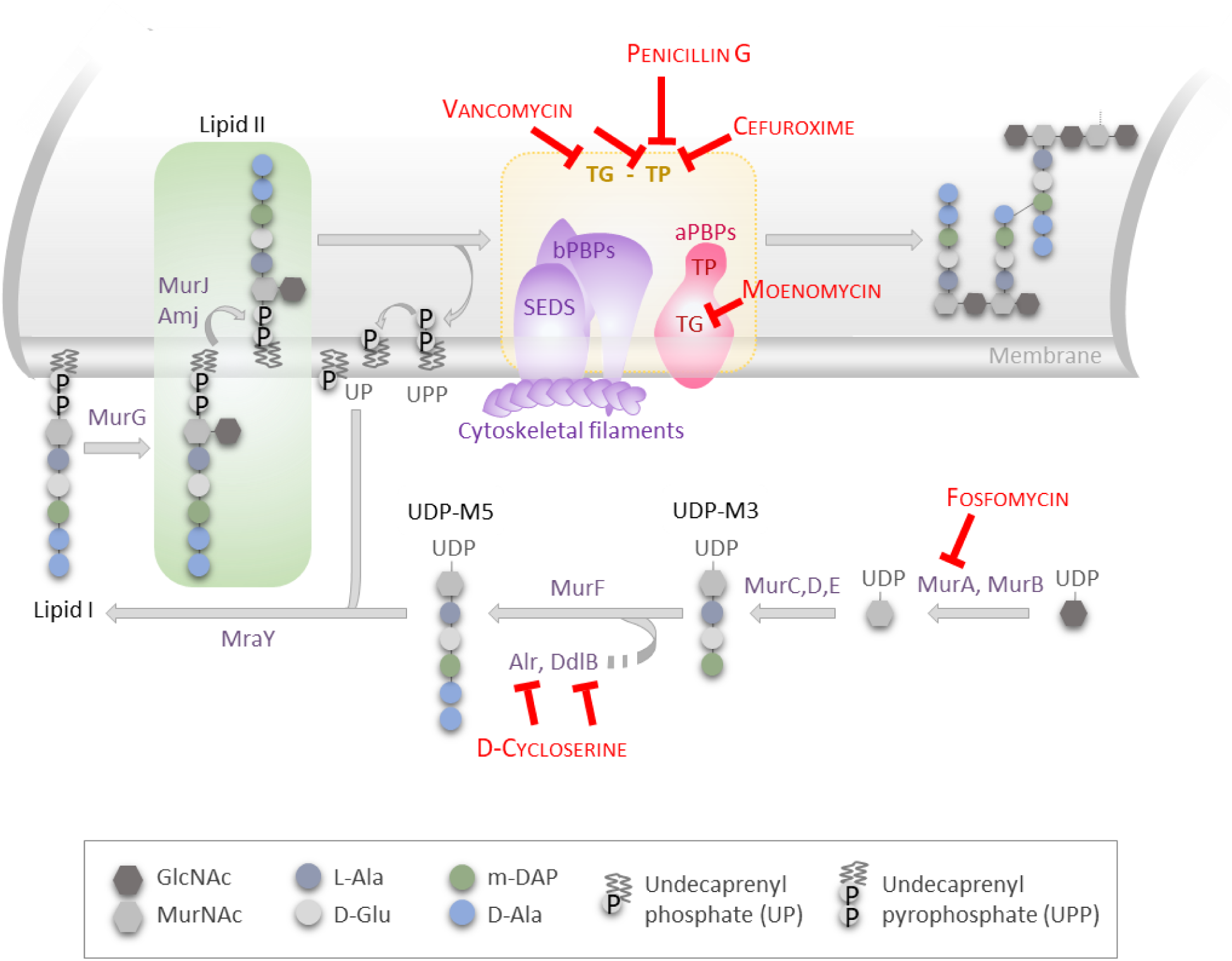
The PG synthesis pathway of *Bacillus subtilis* and antibiotics used in this study. Schematic view of the PG synthesis pathway of *B. subtilis.* The first step of PG synthesis is the formation of UDP-GIcNAc from fructose-6-phospate, which is transformed into UDP-MurNAc by MurA and MurB. L-alanine (L-Ala), D-glutamic acid (D-Glu) and meso-2,6-diaminopimelate (m-DAP) are then added sequentially to the sugar by the action of MurC, MurD and MurE, respectively, to form UDP-MurNAc-tripeptide (UDP-M3). The D-alanine-D-alanine (D-Ala-D-Ala) dipeptide, synthesized by Air and Ddl, is then added by MurF to produce UDP-MurNAc-pentapeptide (UDP-M5). MraY next transfers the soluble UDP-M5 onto undecaprenyl phosphate (UP, also referred as C_55_ lipid carrier), forming the membrane-bound intermediate lipid I in the inner leaflet of the cytoplasmic membrane. Finally, MurG adds a GIcNAc unit, producing lipid II. Lipid II is then flipped to the outer leaflet of the cytoplasmic membrane by the major (MurJ) and the minor (Amj) flippases (Meeske et al. 2015). The externalised PG precursor is linked to a nascent glycan strand by transglycosylation (TG), releasing undecaprenyl-pyrophosphate (UPP) in the process. Concomitantly, crosslink between stem peptides occurs through transpeptidation (TP) reactions (Sauvage et al. 2008). TG and TP reactions are catalysed by the bifunctional aPBPs and by SEDS (shape, elongation, division and sporulation) proteins associated to bPBPs and cytoskeletal proteins. The antibiotics used in this study and their targets are shown in red. Fosfomycin and D-cycloserine inhibit the synthesis of soluble PG precursors in the cytoplasm. Vancomycin, penicillin G, moenomycin and cefuroxime inhibit extracellular TG/TP reactions.

Most rod-shape bacteria use two independent SEDS-containing PG synthesizing machineries to grow (sidewall cylindrical elongation) and to divide (septum formation) (Garde et al. 2021). Sidewall elongation is promoted by the so-called Rod system, which includes the SEDS-type TG RodA (Meeske et al. 2016; Emami et al. 2017) in complex with its cognate bPBPs, and is associated to filaments of the actin homologue MreB (Chastanet and Carballido-Lopez 2012). MreB filaments orient components of the Rod system to move directionally around the cell circumference (Dominguez-Escobar et al. 2011; Garner et al. 2011), promoting oriented insertion of glycan strands in radial hoops (Dion et al. 2019). aPBPs also contribute to sidewall elongation but are thought to act outside the Rod complex, diffusing all over the membrane (Cho et al. 2016) and inserting unoriented glycan strands in a diffuse manner (Dion et al. 2019; Vigouroux et al. 2020). The current model is that the Rod system builds and shapes the main structure of the sacculus while aPBPs add to it to repair it (Cho et al. 2016; Vigouroux et al. 2020; Straume et al. 2020; Murphy et al. 2021). Efficient CW synthesis requires both systems to be functional and thus cooperative interactions must exist between them (Banzhaf et al. 2012; Cho et al. 2016; Cleverley et al. 2019). Furthermore, it has been shown that the balance between the two systems determines the cell diameter in *Bacillus subtilis*: circumferential PG insertion by the Rod complex reduces diameter, while isotropic PG insertion by aPBPs increases it (Dion et al. 2019). *B. subtilis* encodes 4 aPBPs that are not essential nor co-essential, with the major aPBP being PBP1 (encoded by *ponA*) (McPherson and Popham 2003). *B. subtilis* cells lacking *ponA* or all four aPBPs are 20-25% thinner than wild-type cells, while cells become wider upon PBP1 overexpression (Murray et al. 1998; McPherson and Popham 2003; Dion et al. 2019). Conversely, Rod mutants are generally wider than wild-type cells, some losing rod shape and rounding, and cells become thinner as MreBCD expression increases (Chastanet and Carballido-Lopez 2012; Dion et al. 2019). Cell diameter can therefore be used as a proxy to monitor the balance between the activities of the Rod complex and aPBPs in *B. subtilis*.

The activities of the PG biosynthetic systems must be coordinated with the action of PG hydrolases (often called autolysins), which cleave existing bonds in the sacculus to allow CW expansion and incorporation of new glycan strains during growth (Vollmer, Blanot, et al. 2008). Although no PG hydrolase has been found to display MreB-associated circumferential motion, PG hydrolytic activity and in particular the activity of endopeptidases (EPs, which cleave peptide bridges) has been associated to the activity of the Rod system (Domínguez-Cuevas et al. 2013; Sassine et al. 2020). The co-lethal DL-EPs LytE and CwlO have nevertheless been associated to sidewall elongation (Bisicchia et al. 2007; Domínguez-Cuevas et al. 2013; Meisner et al. 2013) and either alone can sustain growth in the absence of the other 41 putative PG hydrolases of *B. subtilis* (Wilson and Garner 2021).

Inhibition of PG assembly by antibiotics weakens the sacculus and often results in cell lysis. Lytic effects are typically thought to result from unbalanced PG synthesis and hydrolysis, in favour of increased PG hydrolytic (autolytic) activity when PG synthesis is inhibited (Tomasz 1979). In agreement with this, PG hydrolytic activity is deregulated in both *ΔmreB* (Tesson et al. 2022) and aPBP mutants (McPherson and Popham 2003). However, the exact crosstalk between PG hydrolases and the PG synthesis machineries, and the specific enzymes among the 42 definite or putative PG hydrolases identified in *B. subtilis* (Smith et al. 2000; Wilson and Garner 2021) that are dysregulated in Rod and aPBP mutants remain unknown. The potentially lethal activity of PG hydrolases is thwarted by multiplicity and functional redundancy, which together with complex regulatory networks hamper efforts to pinpoint their individual effects (Smith et al. 2000; Vollmer, Joris, et al. 2008; Banzhaf et al. 2012). The consequences of inhibition of individual targets of the PG biosynthetic pathway by antibiotics, and how they lead to cell death are also poorly understood. Here, we investigated the effect in growing *B. subtilis* cells of CW antibiotics from two categories: inhibitors of the synthesis of soluble PG precursors in the cytoplasm (fosfomycin and D-cycloserine) and inhibitors of the extracellular TG and TP reactions (vancomycin and penicillin G) (**Fig. 1**). We show that both groups of antibiotics induce growth arrest, thinning of the sacculus and ultimately cell lysis. However, while the lethal activity of vancomycin and penicillin G occurs rapidly and in the absence of any morphological deformation, cells grow for some time following fosfomycin or D-cycloserine addition and widen and bulge, forming large spheres before lysing. We show that PG hydrolytic activity is dysregulated in these cells and accounts for the elevated lysis phenotype of the population but is independent of cell bulging. Visualization of new TP reactions *in vivo* suggested that active PG synthesis occurs at the bulges. By assessing the effect of the CW antibiotics on the localization and dynamics of MreB, as a proxy for the Rod complex, and of PBP1, as well as in Δ*mreB* and Δ*ponA* mutants, we showed that both growth and bulging during depletion of soluble PG precursors depends on PBP1 activity. Altogether, our data show that PG precursors shortage differentially affects the two PG elongation polymerase systems of *B. subtilis*, highlighting differences in their regulation. While Rod system activity is an actively regulated process that depends on the availability of PG precursors, dispersed PG synthesis by the aPBP system is not and continues when PG precursors become limiting, resulting in cell rounding. We suggest that this allows redirection of the available PG precursors to the sites of CW repair to preserve cell integrity while allowing growth to continue during antibiotic threat.

## Results

### Inhibition of soluble PG precursors synthesis induces cell bulging

To visualise the consequences of exposure to CW antibiotics, exponentially-growing *B. subtilis* cells were incubated in the presence of four antibiotics targeting either cytoplasmic or extracellular steps of the PG synthetic pathway: fosfomycin, D-cycloserine, vancomycin and penicillin G. Fosfomycin blocks the first committed cytoplasmic step of PG synthesis by targeting MurA (Aghamali et al. 2019; Dörries et al. 2014; Kahan et al. 1974) and D-cycloserine inhibits synthesis of the D-Ala-D-Ala dipeptide and thus of the soluble PG percursor UDP-MurNAc-pentapeptide (UDP-M5) (Walsh 1989; Batson et al. 2017) (**Fig. 1**). Vancomycin binds to the D-Ala-D-Ala dipeptide of the externalised lipid II and inhibits TG and TP by steric hindrance (Mainardi et al. 2008), and the β-lactam penicillin G mimics the substrate of the TP reaction and thus blocks TP reactions by PBPs (Schneider and Sahl 2010). Chloramphenicol, which targets ribosomes to inhibit protein synthesis (Kitahara et al. 2021), was used as a control antibiotic that does not inhibit PG assembly. In liquid LB medium, chloramphenicol was bacteriostatic and all four CW antibiotics had a bacteriolytic effect on exponentially growing *B. subtilis* (**Fig. S1**). Scanning electron microscopy (SEM) of cells incubated for 2 h in the presence of a lethal concentration of either CW antibiotic showed that penicillin G- and vancomycin-treated cells remained rod-shaped while fosfomycin and D-cycloserine induced strong morphological defects: cells were much wider than untreated control cells and often bulged (**Fig. 2A**). To investigate how these morphological deformations appeared during antibiotic treatment, we acquired time-lapse movies of cells mounted on agarose-coated channel slides. After immobilisation on the slide (t_0_), cells were allowed to grow for 20 min before addition of the antibiotic, and phase contrast and epifluorescence images were collected over 2 h (**Fig. 2B, Fig. S2 and Suppl. movies 1-6**). Cells were semi-automatically segmented using a custom-trained deep-learning model for morphometric analysis, and colony growth and cell morphology were extracted (see **Materials and Methods**). Vancomycin and penicillin G, which can directly access their extracellular target and readily block TG/TP reactions, rapidly (about 10 min after antibiotic addition) arrested growth and induced extensive lysis (**Fig. 2C and Fig. S2, S3)**. Cells lysed without undergoing any morphological deformation, as previously observed for penicillin G in hypotonic NB medium (Kawai et al. 2018). In contrast, the effects of fosfomycin and D-cycloserine on growth appeared later, suggesting that they require the pool of PG precursors to be depleted below a certain threshold. The growth rate of the cell population was nearly unperturbed for some time, then decreased relative to untreated cells (fosfomycin) or the population started lysing (D-cycloserine) while cell diameter gradually increased in the presence of the two antibiotics (**Fig. 2C and Fig. S2, S3**). After 2 h in the presence of fosfomycin, the number of cells normalized by the number of cells at _t0_ (N/N_0_) was reduced of 30 % relative to untreated cells while the surface occupied by them was increased of 190 % (**Fig. S3**), consistent with cell enlargement.

**Fig. 2.**
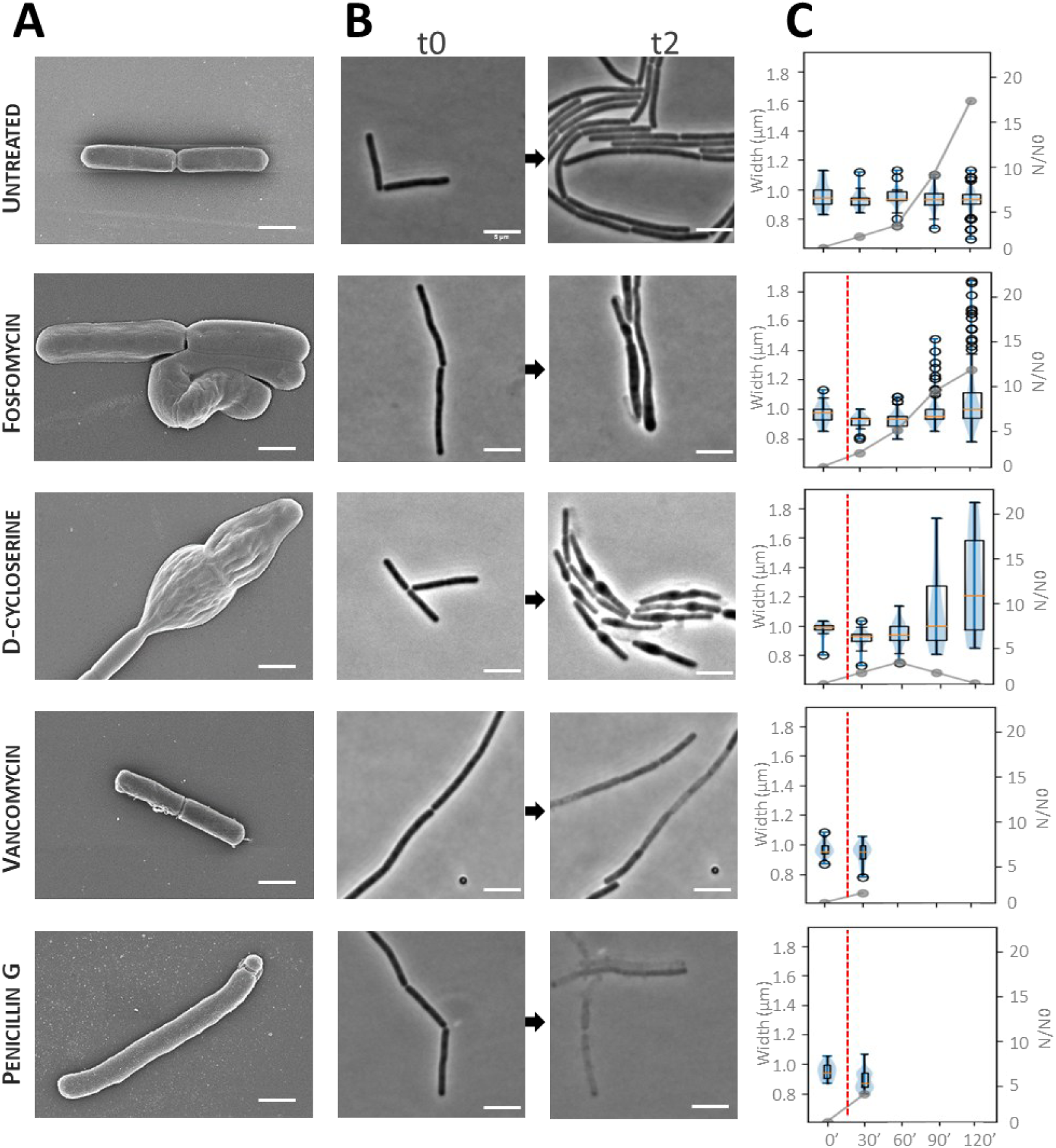
Effect of the CW antibiotics on cell growth and morphology. Cells of the wiId-type *B. subtilis* strain 16S growing in LB medium at 37°C were treated with fosfomycin (25 μg/mL), D-cycloserine (100μg/mL), vancomycin (2μg/mL) or penicillin G (20μg/mL). **A.** Representative SEM images. Cell were grown to exponential phase before addition of the antibiotic, and incubated 2 h in presence of antibiotic before sampling. Scale bar, 1 μm. **B.** Phase contrast microscopy images before (t0) and 2 h after (t2) addition of the antibiotics. Exponentially growing cells were stained with 10 μg/mL NileRed as membrane dye and spotted on agarose pads prepared in LB with 10 μg/mL NileRed (t0). Antibiotics were added after allowing the cells to settle on the pad for 20 min. Phase contrast and epifluorescence (membrane staining) images were taken every 2 min (see accompanying Fig. S2 and corresponding Suppl. movies S1-S6). Scale bar, 5 μm. **C.** Quantification of the colony growth and of the cell diameter from time-lapses microscopy movies. Exponentially growing cells were stained with 10 μg/mL NileRed as membrane dye and spotted on agarose pads prepared in LB with 10 μg/mL NileRed (t0). Antibiotics were added after allowing the cells to settle on the pad for 20 min (dashed line). The evolution of the number of cells (N), normalized by the number of cells at t0 (N0) is shown as grey curves, with the corresponding axis on the right. The curves are a mean of 3 independent experiments. Cell width distributions (in μm) are depicted as boxplots, with the corresponding axis on the left.

### Antibiotic-induced cell bulging is partially dependent on deregulated PG hydrolytic activity

During fosfomycin and D-cycloserine treatment, significant lysis was observed at the single cell level as surviving cells continued to enlarge (**Fig. S2)**. This elevated lysis phenotype may contribute to the reduced growth rate of the population. Noteworthy, bulging (generally considered to precede CW antibiotic-induced cell lysis) and cell lysis were not correlated. Lysis was indistinctly observed for bulged and seemingly unaffected rod-shaped cells (**Fig. S2 and S4A**), indicating that it is not a consequence of bulging, and that it may result from deregulated autolytic activity instead. We previously showed that in *B. subtilis* millimolar concentrations of Mg^2+^ in the growth medium suppress the enhanced lysis and the morphological defects of Δ*mreB* mutants through their inhibitory effect on deregulated PG hydrolases (Dajkovic et al. 2017; Tesson et al. 2022). Addition of 20 mM Mg^2+^ rescued growth and the elevated lysis phenotype of fosfomycin- and D-cycloserine-treated cells (**Fig. S1 and S4B**), indicating unbalanced PG hydrolytic activity in these cells. However, high Mg^2+^ did not rescue cell bulging, which became still more pronounced and led to large spherical cells (**Fig. 3A and Suppl. movie 7**). In agreement with this, mutants of *cwlO* and *LytE,* which encode the two synthetically lethal EPs of *B. subtilis* required for cell growth (Bisicchia et al. 2007; Wilson and Garner 2021), also bulged in the presence of D-cycloserine (**Fig. S5B**, **Suppl. movie 8, 9, 10, 11**). Interestingly, lysis in the presence of D-cycloserine was reduced in a *ΔlytE* mutant (**Fig. S5**), suggesting that LytE activity is deregulated in these conditions. LytE was recently shown to be inhibited by Mg^2+^ (Wilson and Garner 2021). However, lysis of the *ΔlytE* mutant was further reduced in the presence of Mg^2+^, indicating that other PG hydrolases also inhibited by Mg^2+^ and deregulated in the presence of D-cycloserine. In a *ΔlytABCDE* mutant, D-cycloserine-induced lysis was further reduced but still inhibited by Mg and bulging was almost not observed, especially in the presence of magnesium **(Suppl. movie 12, 13)** (Kawai et al. 2023). These results suggest that antibiotic-induced inhibition of soluble PG precursors synthesis induces both dysregulated PG hydrolytic activity and cell bulging *in B. subtilis*, and that these are partially dependent of each other.

**Fig. 3.**
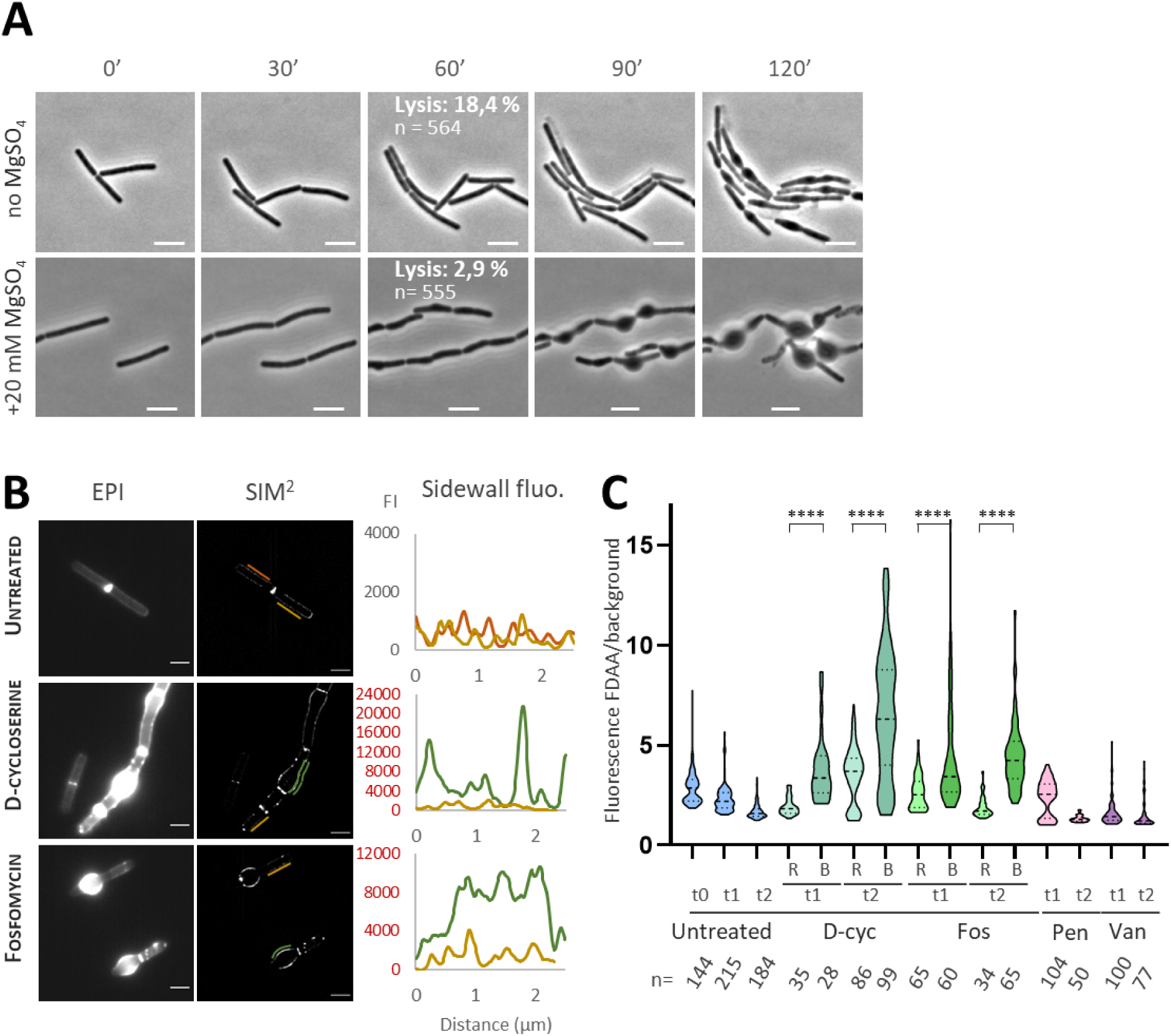
Cell bulging is partially dependent on deregulated PG hydrolytic activity, FDAA labelling revealed active PG synthesis in bulges. Antibiotics were used at the following concentrations: 100 μg/mL D-cycloserine (D-cyc), 25 μg/mL fosfomycin (Fos), 20 μg/mL penicillin G (Pen) or 2 μg/mL vancomycin (Van). **A.** Cell were grown in tube in LB +/- 20 mM MgsO_4_ until exponential phase before being spotted on agarose pad prepared in the same medium. Timelapse on agarose pad were realised in LB (upper row) and in LB additionned with 20 mM MgSO_4_ (lower row). Images were taken every 2 min, 1 image every 30 min is presented here (scale bar: 5μm). D-cycloserine was added at t0 (0’) and used at 100 μg/mL. The percentage of lysis at 60 min is indicated in white on the 60’ image. See corresponding Suppl. movies S4, S7. Scale bar: 5μm. **B.** Epifluorescence (same settings for all conditions) and SIM^2^ images of sBADA labelling of *B. subtilis* wt cells at 1 h (t1) after addition of antibiotic (scale bar: 2 μm). The intensity profiles (fluorescence intensity Fl along the sidewall) are shown for two cells of each condition. **C.** Quantification of sBADA fluorescence intensity measured along the sidewall and normalized by the background. For D-cycloserine and fosfomycin treated cells, rod-shaped (“R”) and bulging (“B”) cells are shown. Significant difference between rod and bulging cells is represented as “*’’.

### PG synthetic activity occurs at the bulges of cells treated with inhibitors of PG precursors synthesis

Under the experimental conditions used here, the pool of soluble PG precursors is virtually depleted after 2h incubation with fosfomycin, and UDP-M3 greatly accummulates in D-cycloserin-treated cells, while soluble PG precursors accummulate in the presence of vancomycin and penicillin G (Cornilleau et al. 2023), confirming that PG precursor synthesis is blocked. We hypothesized that bulging in the presence of fosfomycin and D-cycloserine might result from a weaken CW in cells that continue to expand their sacculi normally but insert material at a severely reduced rate. Sacculi might then become thinner and/or less dense, and reach a threshold below which they are unable to counteract the high internal pressure and maintain rod shape, leading to cell bulging and rounding. In this scenario, cells treated with vancomycin or penicillin G might stop growing and lyse before the thickness or density of their sacculi reach such threshold, allowing rod shape to be maintained. Using UPLC analysis, we recently showed that after 2h treatment, the fraction of PG cross-links is reduced by about 25 % - 12 % fold by vancomycin and penicillin G, as expected upon inhibition of TP reactions, but is almost unaffected (reduced by 5 % and 0 %) by fosfomycin and D-cycloserine (Cornilleau et al. 2023). We then measured the thickness of the CW on transmission electron microscopy (TEM) images and found it to be significantly reduced by all antibiotics (**Fig. S6A**). The most pronounced effect was produced by fosfomycin and vancomycin, which both thinned the sacculi down to 56 % relative to untreated cells. Thus, no correlation was found between the thickness or the fraction of cross-links of the CW and the morphology of cells incubated in the presence of either group of PG synthesis inhibitors. We concluded that CW thinning or reduced crosslinking are not sufficient to induce cell bulging. Further inspection of the TEM images showed that the sacculi of cells treated with all four antibiotics had holes and ragged material, with fragments ‘peeling off’ (**Fig. S6C**), consistent with the observed lysis and dysregulated PG hydrolytic activity. Interestingly, CW abnormalities with thick PG deposition points, were noticeable in cells treated with D-cycloserine but not in cells treated with the TG/TP inhibitors (**Fig. S6D**).

We then wondered if bulging could result from abnormal PG synthetic activity. To investigate this, we monitored PG synthesis using the green-emission fluorescent D-amino acid (FDAA) probe sBADA (sulfonated BODIPY-FL 3-amino-d-alanine) (Hsu et al. 2017) (**Fig. S7A**). In *B. subtilis,* FDAAs are incorporated in the 5^th^ position of the stem peptide by D,D transpeptidation, catalysed by PBPs (Kuru et al. 2012, 2019). Thus, FDAAs labelling is typically used to report PG cross-linking activity as a proxy of PG biosynthesis *in vivo*. Furthermore, incorporated FDAAs are cleaved by the DD-carboxypeptidase PBP5 (encoded by *dacA*), which removes the C-terminal D-Ala residue from the PG precursor pentapeptide chain (Kuru et al. 2012), and thus can also reflect PG turnover *in vivo*. In untreated cells, sBADA labelling was visible at the septa and along the cylindrical sidewalls and dimmer at the poles (**Fig. 3B**, **Fig. S7B, C, D**), as previously reported using other FDAAs (Dajkovic et al. 2017; Tesson et al. 2022). To improve optical sectioning and increase lateral resolution we used lattice-SIM (structural illumination microscopy) coupled to the SIM² image reconstruction algorithm (Chen et al. 2023). Using SIM² we observed that sBADA fluorescence was distributed in a regular discrete pattern along the sidewalls, consistent with the sites of active Rod complex activity in growing cells. In cells treated with vancomycin or penicillin G, sBADA labelling became progressively faint and virtually disappeared (**Fig. 3C**, **Fig. S7A, C, D**), consistent with the inhibition of TP reactions. However, the average sBADA signal was much more intense in cells incubated with D-cycloserine or fosfomycin than in untreated cells after either 1h or 2h incubation (**Fig. 3B, C and Fig. S7A, C, D**), in both the wild-type and the *ΔdacA* background (**Fig. S8**). Notably, the very intense sBADA labelling was coming from bulging cells, while rod-shaped cells in the same fields were less stained (**Fig. 3B, C, Fig. S7C, D**). The pattern of fluorescence also appeared less regular in bulged cells than in untreated cells (**Fig. 3B, Fig. S7C, D**). Taken together, these findings indicate that cell bulging in the presence of PG precursors synthesis inhibitors is associated with increased and more irregular TP activity.

### Rod complex activity halts and PBP1 diffusion slows down when soluble PG precursors are depleted

The high TP activity in the bulges of D-cycloserine- and fosfomycin-treated cells could result from Rod complex and/or aPBPs PG synthetic activity at these sites when the pool of PG precursors is being depleted. MreB circumferential motion depends on active PG synthesis and is believed to reflect PG synthesis by the Rod complex (Dominguez-Escobar et al. 2011; Garner et al. 2011; Rueff et al. 2014; Schirner et al. 2015; Meeske et al. 2016). Furthermore, the levels of lipid-linked PG precursors regulate MreB membrane association (Schirner et al. 2015; Sun et al. 2023). In agreement with this, addition of either D-cycloserine or fosfomycin rapidly stopped MreB directed motion and progressively decreased total MreB filament density in the membrane (**Fig. 4A-B and Fig S9A**). The same effect was observed when we monitored Mbl (**Fig S9B**), the other isoform of MreB, essential in *B. subtilis* and also associated to the Rod complex (Carballido-López et al. 2006; Dominguez-Escobar et al. 2011). Vancomycin and penicillin G drastically reduced MreB directed motion as well but did not cause MreB filaments to disassemble from the membrane (**Fig S9A**) (Dominguez-Escobar et al. 2011; Garner et al. 2011). We next imaged a functional mNeonGreen-PBP1 fusion (Cho et al. 2016) in treated and untreated cells. Two dynamic populations of PBP1 in *B. subtilis* (Cho et al. 2016) and of PBP1a and PBP1b *in E. coli* (Lee et al. 2016; Vigouroux et al. 2020) were previously reported to co-exist in the cell membrane: an immobile or ‘slow’ population with virtually zero diffusion coefficient thought to monitor sites of PBP1 activity and a diffusive ‘fast’ population thought to search for new sites of insertion, presumably CW damage defects (Cho et al. 2016). Analysis of single-particle trajectories in untreated cells using cumulative distribution functions (CDFs) confirmed the two states and gave diffusion coefficients (D) of PBP1 similar to those reported previously (D diffusive = 2.10^-3^ µm^2^ s^−1^ and D immobile = 10^-4^ µm^2^ s^−1^) (Cho et al. 2016). Within 60 min in the presence of fosfomycin or D-cycloserine, PBP1 signal was enriched at bulging sites (**Fig. 4C**) and the immobile fraction of PBP1 slightly increased and the diffusion coefficient D1 decreased (**Fig. 4D, E, F and Fig. S9C, D**). However, in the presence of vancomycin or penicillin G the dynamic behaviour of PBP1 was strongly affected: the immobile fraction drastically increased, and the diffusion coefficient were drastically reduced (**Fig S9C, D**). These results suggested that PBP1 dynamic populations are mostly unaffected during fosfomycin / cycloserine treatment but highly perturbed when the cells are exposed to vancomycin and penicillin G.

Taken together, these results suggest that inhibition of PG precursors synthesis rapidly arrests Rod complex activity but not aPBPs activity, which may continue at low PG precursor levels and, associated with deregulated autolysis activity, may induce cell bulging.

**Fig. 4.**
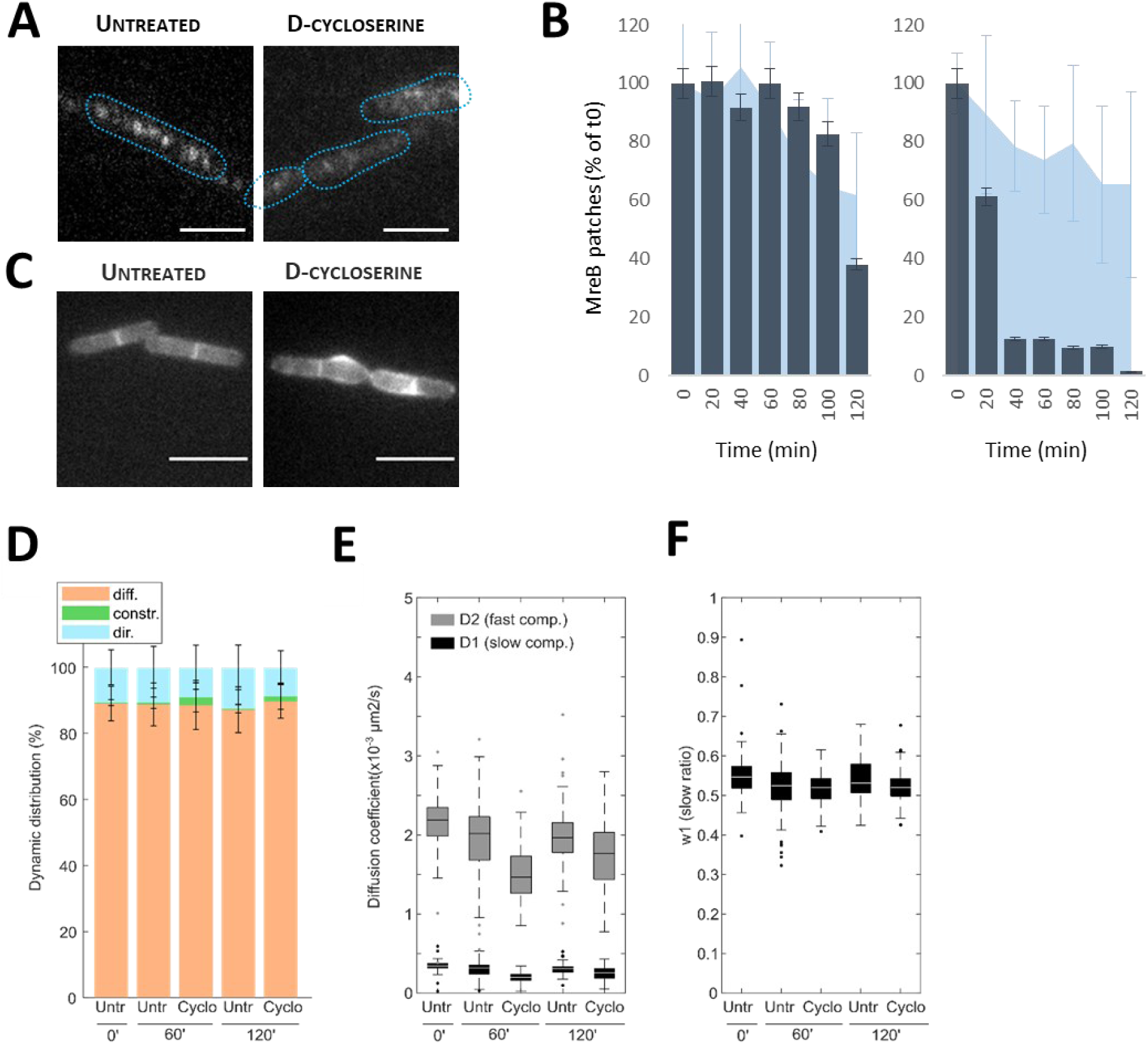
Inhibitors of PG synthesis affect MreB, not PBP1. **A.** GFP-MreB patches visualisation in TIRF microscopy after 1 h treatment with 100 μg/mL D-cycloserine (scale bar: 2μm). The cells outline are shown as dotted line. **B.** Dynamic behaviour and density of MreB patches for an untreated control and for cells treated with 100 μg/mL D-cycloserine. Cells were grown until exponential phase (t0) before addition of CW antibiotic. GFP-MreB was imaged in TIRF microscopy at t0 and every 20 min after. The dynamic behaviour of MreB patches is based on MSD analysis, only the percentage of directed patches (normalized by the percentage at t0) is represented here (dark blue histograms). The density of MreB patches is represented as a percentage of the density at t0 (light blue area). **C.** PBP1 localisation. *P*_hyperspank_*-mNeonGreen-ponA* strain untreated and treated with 100 μg/mL D-cycloserine during 1 h before imaging in epifluorescence (scale bar: 5 μm). **D.** PBP1 dynamic behaviour. *P*_hyperspank_*-mNeonGreen-ponA* strain was grown until exponential phase (t0) before addition of CW antibiotic (D-cycloserine 100 μg/mL, fosfomycin 25 μg/mL) and furter grown during 1 h and 2 h before imaging. **E.** PBP1 diffusion was quantified at single cell levels (or filament levels if impossible to resolve) using CDF analysis with two components D1 and D2, resp. slow and fast components **F.** w1 is the weight of slow component in the population.

### Growth and bulging of D-cycloserine-treated cells depend on PBP1 activity

To investigate if aPBP activity was responsible for cell growth and bulging when PG precursors are limiting, we looked at the effect of moenomycin on D-cycloserine-treated cells. Moenomycin A is a natural phosphoglycolipid, antibiotic that specifically targets aPBPs (Van Heijenoort et al. 1978; Ostash and Walker 2010) (**Fig. 1**). However, moenomycin does not inhibit the activity of the transglycosylases FtsW and RodA (Meeske et al. 2016; Zhao et al. 2017; Emami et al. 2017). *B. subtilis* can grow in the absence of aPBPs (McPherson and Popham 2003) by inducing the Rod system (Emami et al. 2017), and thus is intrinsically resistant to moenomycin (Meeske et al. 2016; Zhao et al. 2017; Emami et al. 2017). In agreement with this, growth of wild-type cells was unaffected by the addition of moenomycin (**Fig. 5A**). However, in the presence of a sub-inhibitory concentration of D-cycloserine, growth was completely inhibited by moenomycin (**Fig. 5A**). These results indicated that aPBPs activity is needed for growth when synthesis of PG precursors is perturbed. To confirm this, we looked at the effect of D-cycloserine in a Δ*ponA* null mutant. In contrast to wild-type cells, Δ*ponA* cells did not grow in the presence of the sub-inhibitory concentration of D-cycloserine (**Fig. 5A**). We next monitored the growth and morphology of the mutant in time-lapse experiments (**Suppl. movies 14-15**). Untreated Δ*ponA* cells exhibited the typical reduced diameter (Popham and Setlow 1995; McPherson and Popham 2003; Dion et al. 2019) and reduced growth rate of the population relative to wild-type cells (**Fig. S10A-B**). Upon addition of D-cycloserine, growth of Δ*ponA* cells was arrested and the bulging phenotype observed in wild-type cells was suppressed (**Fig. 5B, Fig. S10A, B**). D-cycloserine-treated cells did not bulge either when aPBPs were inhibited by moenomycin (**Fig. S11A**). These findings confirmed that the major aPBP, PBP1, is required for both growth and cell bulging in the presence of PG precursor inhibitors. The diameter of *B. subtilis* is determined by the balance between the widening effect of aPBPs and the thinning effect of the Rod system (Dion et al. 2019). When Δ*mreB* mutant cells were grown in the presence of D-cycloserine, their bloated phenotype was more severe than in the absence of the antibiotic and they became spherical (**Fig. 5B, Fig. S10 A-B, Suppl. movies 16-17**). This was expected because Mbl is still active in Δ*mreB* cells and can support some rod-shaped morphology (**Fig. S10A**) (Kawai, Asai, et al. 2009) but D-cycloserine induced disassembly of both MreB (**Fig 4A-B**) and Mbl (**Fig S9B**) filaments. We concluded that bulging of cells in the presence of antibiotics that deplete the pool of PG precursors results from disassembly of the Rod complex (**Fig. 4B**) while aPBP activity may continue diffuse insertion of PG around the cell. To confirm this, we monitored sBADA incorporation along the sidewalls of D-cycloserine-treated cells in which aPBP activity was inhibited. As expected, in the presence of D-cycloserine, the increased sBADA labelling observed in wild-type cells was strongly reduced both in cells lacking *ponA* and in cells treated simultaneously with moenomycin (**Fig. 5C and Fig. S11A-C**). In contrast, the staining of Δ*mreB* cells was enhanced both at the division sites and along the sidewalls when the cells were treated with D-cycloserine (**Fig. 5C and Fig. S11C**).

**Fig. 5.**
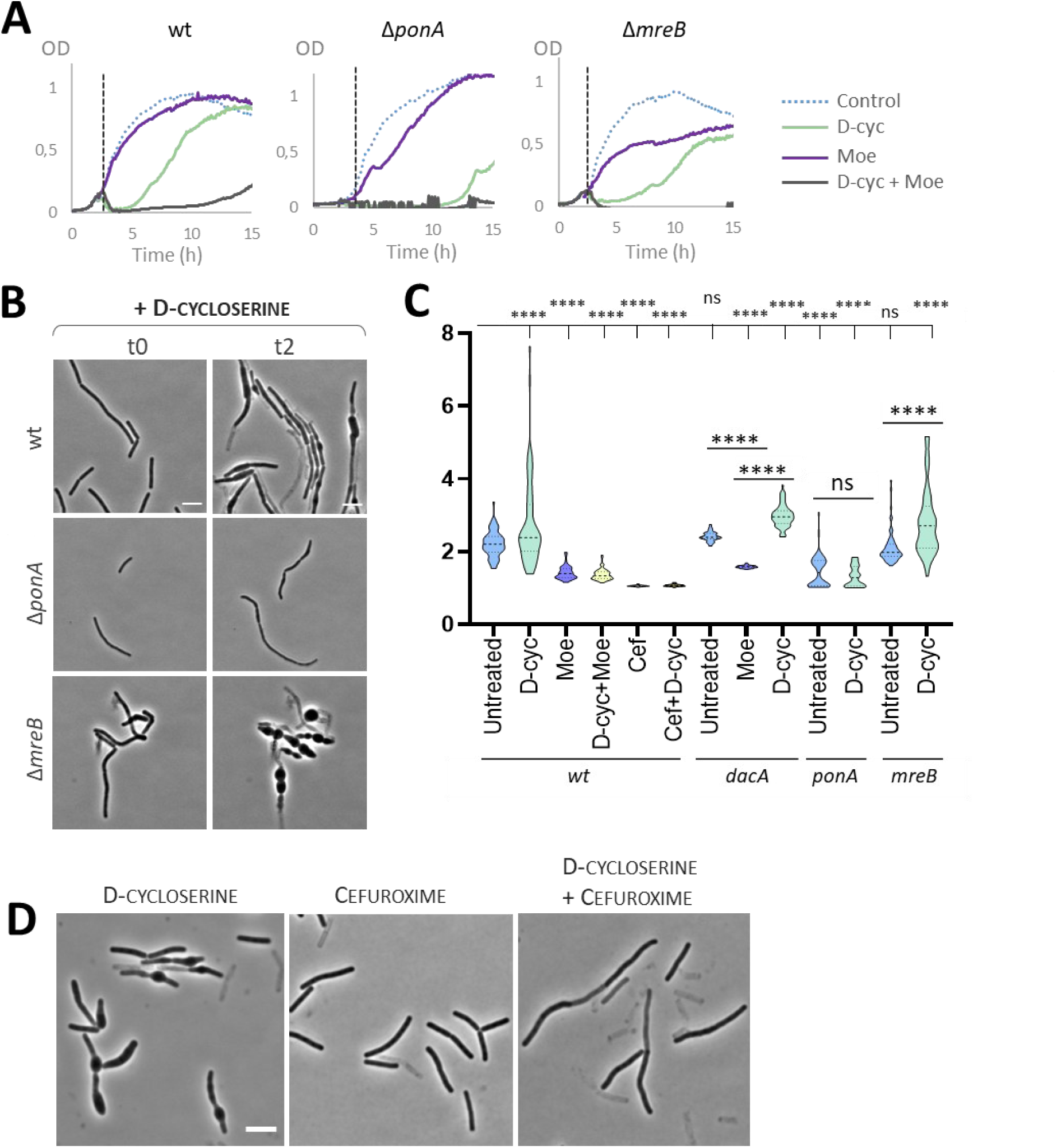
PBP1, encoded by ponA, is likely to be responsible for the remaining PG synthesis in D-cyloserîne treated cells and thus for the deformations of these cells. **A.** Growth curves of the wt strain, *ΔponA* and *ΔmreB* mutants with addition of antibiotic in exponential phase of growth (grey dashed line). Antibiotics were used at the following concentrations: 20 μg/mL D-cycloserine (D-cyc), 50 μg/mL moenomycin (Moe). **B.** Phase contrast images of the wt strain, *ΔponA* and *ΔmreB* mutants treated with 100 μg/mL D-cycloserine (scale bar: 5 μm). Cell were grown in LB medium until exponential phase (t0) before addition of antibiotic, and incubated 2 h (t2) in presence of antibiotic before sampling. See corresponding **Suppl. moviesS14-S17**. **C.** Quantification of sBADA fluorescence intensity measured along the sidewall and normalized by the background for the wt strain, *ΔdacA, ΔponA* and *ΔmreB* mutants untreated and treated with antibiotics. Significance in represented as “*”; n>55. Antibiotics were used at the following concentrations: 100 μg/mL D-cycloserine (D-cyc), 50 μg/mL moenomycin (Moe), 5 μg/mL penicillin G (Pen), 0.05 μg/mL cefuroxime (Cef). **D.** Phase contrast images of the wt strain treated with 100 μg/mL D-cycloserine, 0,05 μg/mL cefuroxime or both antibiotics (scale bar: 5 μm).

Noteworthy, labelling of both Δ*ponA* and moenomycin-treated cells was reduced to a similar extent (about 35-40%) in the presence and in the absence of D-cycloserine (**Fig. 5C and Fig. S11A, B, C**). This suggested either higher DD-carboxypeptidase activity when aPBP function is inhibited or, alternatively, that aPBP TP activity has an important contribution to sBADA incorporation in exponentially growing *B. subtilis* cells. To discriminate between these two hypotheses, we monitored sBADA incorporation in the Δ*dacA* mutant treated with moenomycin (**Fig. S11D**). Inhibition of aPBP activity by moenomycin also reduced the sBADA labelling of the Δ*dacA* mutant of about 35%, like in wild-type cells (**Fig. 5C and Fig. S11A-D**), indicating that DD-carboxypeptidase activity is not increased in response to the loss of aPBP activity. Thus, the sBADA signal reduction measured in Δ*ponA* cells relative to wild-type cells reflects the TP activity of PBP1. In agreement with this, the degree of crosslinking has been reported to be reduced of a similar order of magnitude (20%-24%) in a *ΔponA* mutant (Atrih et al. 1999; Cornilleau et al. 2023).

### The TP activity of PBP1 is needed for growth and bulging when PG precursors are limiting

Our experiments above indicated that PBP1 activity is required for growth and bulging during reduced PG precursor availability. Moenomycin inhibits the TG activity of bifunctional aPBPs by competing with the growing chain of PG in the aPBP active site (Sauvage et al. 2008; Ostash and Walker 2010). However, although the TG and TP domains of aPBPs are distinct, inhibition of TG by moenomycin or by mutation affects as well the TP activity of aPBPs and completely blocks PG polymerisation (Sauvage et al. 2008). Thus, our moenomycin experiments could not reliably inform on whether it is the TG or the TP activity of PBP1, or both, that promote cell bulging. In contrast, the TG of aPBPs can proceed while the TP domain is inhibited by antibiotics, mutation or is completely deleted (Terrak et al. 1999; Born et al. 2006). Selective antibiotics allow investigation of the enzymatic activity of individual PBPs (Sharifzadeh et al. 2020). We thus turned to cefuroxime, a β-lactam reported to preferentially inhibit the TP activity of PBP1 over bPBPs in *B. subtilis* (Sassine et al. 2017; Sharifzadeh et al. 2020) while penicillin G preferentially inhibits bPBPs. Consistently, the Δ*ponA* mutant was found to be more resistant to cefuroxime than the wild-type (Patel et al. 2020). When we treated wild-type cells with sub-inhibitory concentrations of D-cycloserine and cefuroxime simultaneously, growth and bulging were inhibited (**Fig. 5D**), and sBADA staining along the sidewalls was strongly reduced (**Fig. 5C**). Taken together, these results suggest that the TP activity of PBP1 is required for growth and bulging during reduced PG precursor availability.

## Discussion

### Blocking the synthesis of soluble precursors provokes irregularities in the CW and cell bulging

Cell imaging in phase contrast and SEM microscopy showed that CW antibiotics provoked different consequences depending on their target: drugs inhibiting the synthesis of PG precursors triggered cell deformation, with or without lysis, whereas in the presence of TG/TP inhibitors cells lysed without prior deformation. Cell deformations seemed to be linked with growth and active PG synthesis (or at least an active TP), as visualized by FDAA incorporation.

### Cell bulging results from PG synthesis by aPBPs

In *B. subtilis* two PG elongation machineries co-exist: aPBPs and the Rod complex. We monitored both machineries by tracking PBP1 (the main aPBP), and MreB as a proxy of the Rod complex. It has been proven that the Rod complex activity depends on PG synthesis and is compromised by several antibiotics (Dominguez-Escobar et al. 2011; van Teeffelen et al. 2011; Schirner et al. 2015; Meeske et al. 2016). By monitoring MreB dynamics, we confirmed that the Rod complex activity is dramatically affected by CW antibiotic treatment, MreB patches being stopped by D-cycloserine, fosfomycin, penicillin G and vancomycin. This has two consequences on PG synthesis: an arrest of PG synthesis by the Rod complex and unbalanced activity of aPBPs. Indeed, MreB has been proven to be necessary for PBP1 proper localization (Kawai, Daniel, et al. 2009).

We then investigated the effect of CW antibiotics on aPBPs. Vancomycin triggered a mislocalization and a diminution of PBP1 diffusion coefficient, traducing a huge effect of this antibiotic on aPBPs activity. The effect of TG/TP inhibitors on aPBPs has already been shown in *E. coli*, where β-lactam treatment (inhibiting TP) slowed-down PBP1A diffusion, effect associated with a diminution of PBP1A activity (Lee et al. 2016). However, in D-cycloserine-treated cells, PBP1 localisation was similar than in the untreated control cells, at the exception of an enrichment in the bulges of deformed cells. This enrichment in PBP1 correlated with intense PG synthesis as monitored by FDAA staining. A deletion of *ponA* (coding for PBP1) or a treatment with moenomycin (inhibiting TG activity of aPBPs) prevented FDAA incorporation and cell bulging during D-cycloserine treatment.

Altogether, these results indicate that while the activity of the Rod complex is arrested by the PG antibiotics tested, PBP1 remains active in D-cycloserine-treated cells and may be responsible of their bulging (resulting from PBP1-dependent diffuse insertion of PG). This is in good agreement with a recent study showing that PBP1 can work at lower levels of PG precursors than the Rod complex (Kawai et al. 2023).

Changes in the PG features produced by CW antibiotics (Cornilleau et al. 2023), show that a 2h treatment with inhibitors of PG precursors synthesis (bacitracin, fosfomycin, D-cycloserine and tunicamycin) increases the proportion of trimers of the network. aPBPs being suspected to realized PG synthesis in these conditions (this study), they might be responsible for this increase in trimers, which is consistent with the reported decrease in trimers in a *ponA* mutant (Atrih et al. 1999). Trimers interconnect multiple strands of PG and have been suggested to help strengthening the PG meshwork, especially when glycan strands are short (de Pedro and Cava 2015). This suggest that either aPBPs produce shorter PG strands that should be crosslinked as trimers to ensure CW integrity, or that their TP activity tends to crosslink more trimers than the TP activity of bPBPs. This is in good agreement with the current model proposing that aPBPs are filling gaps in the CW and repairing the PG network.

### Proposed model: differential action of PG synthesis inhibitors on aPBPs and the Rod complex

Taken together, the results reported here allow us to propose a model to explain the chemical and morphological perturbations observed after CW antibiotic treatment (**Fig. 6**). During normal growth, PG is synthetized along the sidewall by two elongation systems: the Rod complex inserts PG chains circumferentially to ensure elongation of the cell while aPBPs fill in the gaps in the network by inserting PG in a diffusive manner. The balance between the two systems ensures the maintenance of rod shape and a constant diameter (Emami et al. 2017; Straume et al. 2020; Dion et al. 2019). TG/TP inhibitors, by inhibiting polymerisation of PG, provoke an arrest of both systems: PG precursors accumulate, cells quickly stop growing and lyse. Inhibitors of PG precursors synthesis, in contrast, provoke a dissolution of Rod complexes while aPBPs remain active. aPBPs keep working, ensuring cell survival but with a larger width and ultimately leading to cell rounding as a consequence of diffusive PG insertion. This suggests that, unlike the Rod complex, aPBPs are not sensing perturbations in PG precursors pathway.

**Fig. 6.**
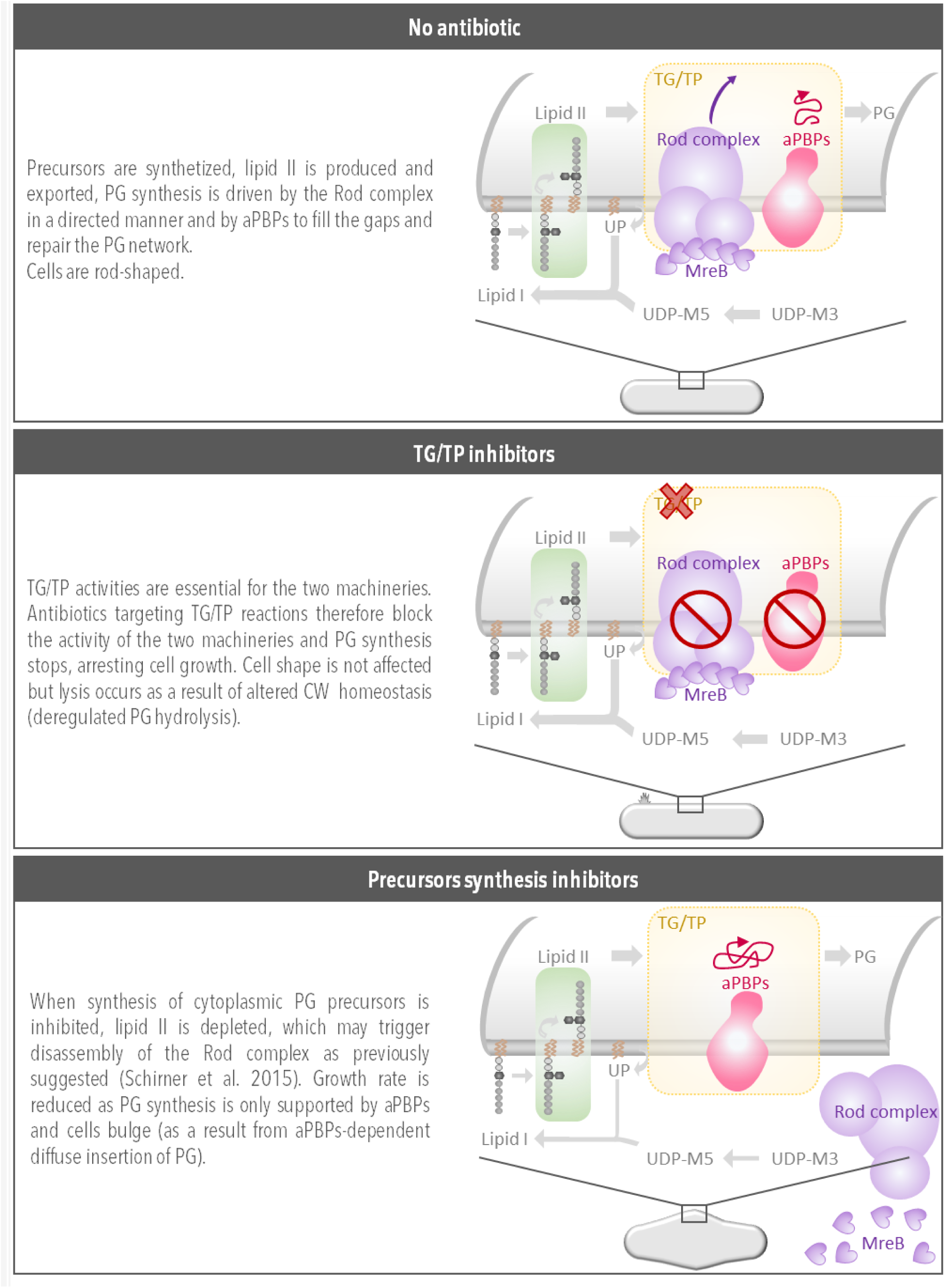
Proposed model explaining the chemical and morphological perturbations observed after cell-wall antibiotic treatment.

aPBPs may respond to the metabolic state of the cell. It has been shown that deletion of genes coding for enzymes involved in different metabolic pathways results in thinner cells (Juillot et al. 2021). In *B. subtilis*, cell diameter is determined by the balance between activities of the Rod complex, which reduces it, and aPBPs, which increase it (Dion et al. 2019). Thus, a thinner diameter in the metabolic mutants might correlate with a decrease of aPBPs activity. This suggests that while the activity of the Rod complex relies on the flow of precursors in the PG synthesis pathway, the activity of aPBPs responds to the overall metabolic state of the cell.

Another interesting idea recently proposed is that aPBPs may be able to physically feel the holes in the CW through a long intrinsically disordered region that scans the PG network (Brunet et al. 2022). Detection of a gap in the PG network would then localize aPBPs, enabling them to synthetize PG at the right place to repair it.

## Conclusion

Inhibitors of TG/TP and inhibitors of PG precursors synthesis provoke different defects: with the former cells lyse without deformation whereas with the latter the control of cell diameter is lost, leading to cell bulging and widening. These morphological defects seem to be due to an anisotropic PG insertion by aPBPs – the Rod complex being stopped. This suggest that aPBPs and the Rod complex are differentially regulated and respond to different stimuli.

All these observations give us insight about PG synthesis and about CW antibiotics mode of action. However, how cells are dying is still to discover. How TG/TP inhibitors trigger lysis so quickly? Are autolysins at play? Cell deformation and lysis appeared uncoupled after treatment by inhibitors of PG precursors synthesis; what is deciding for the faith of a cell?

## Materials and Methods

### Strains and media

Cells were grown in LB medium at 30°C for overnight cultures and at 37°C for day cultures.

We used *B. subtilis* 168 trpC2 as wild type (*wt*) strain. Δ*mreB*(neo) strain is RCL413 from (Billaudeau et al. 2019), Δ*ponA*::spc is PS2062 from (Popham and Setlow 1995), mNeonGreen-PBP1 is MK0287 from (Cho et al. 2016), Δ*dacA*::spc is from (Juillot et al. 2021).

*gfp-mreB* RCL421 strain was constructed in two steps. First, upstream *mreB* region containing neomycin resistance cassette was amplified from strain RCL414 with primers cc181/cc186 and *gfp-mreB* was amplified from strain RCL238 (Billaudeau et al. 2017) with cc182/cc185; fragments were assembled by isothermal assembly and the subsequent product was transformed into *Bacillus subtilis* 168 trpC2 producing strain RCL420. In a second time, GFP was monomerized (A206K mutation): DNA from strain RCL420 was amplified with primers cc181/AC1282 and with cc182/AC1281. Fragments were assembled by isothermal assembly, the resulting product amplified by cc181/cc182 and transformed into *Bacillus subtilis* 168 trpC2 producing strain RCL421.

#### Primers

cc181- GTCATGGGCCTTCCTATATC cc182- GGATGTGCTCCAGTGCTTTC

cc185- GATACATACATTTAAGGAGGTAAACTAATGAGTAAAGGAGAAGAACTTTTCACTG

cc186- TAGTTTACCTCCTTAAATGTATGTATCTTCCTTTCTTAAAGC

AC1281- CAATCTAAACTTTCGAAAGATCCCAACG

AC1282- CGAAAGTTTAGATTGTGTGGACAGGTAATG

### Antibiotics

Stock solution of antibiotics were prepared as follow: fosfomycin 10 mg/mL in H_2_O, D-cycloserine 10 mg/mL in H_2_O, moenomycin 10 mg/mL in DMSO, penicillin G 10 mg/mL in H_2_O, vancomycin 1 mg/mL in H_2O_, lysozyme 10 mg/mL in H_2_O, chloramphenicol 10 mg/mL in ethanol, cefuroxime 1 mg/mL in DMSO.

### Concentrations used in this study

❯ SEM experiments: fosfomycin 25 µg/mL, D-cycloserine 20 µg/mL, vancomycin 2 µg/mL, penicillin G 20 µg/mL.
❯ TEM experiments: fosfomycin 5 & 25µg/mL, D-cycloserine 20 & 100 µg/mL, vancomycin 0.25 & 2 µg/mL, penicillin G 5 & 20 µg/mL chloramphenicol 2 µg/mL.
❯ Timelapses: fosfomycin 25 µg/mL, D-cycloserine 100 µg/mL, vancomycin 2 µg/mL, penicillin G 20 µg/mL, lysozyme 20 µg/mL, chloramphenicol 5 µg/mL.
❯ FDAA staining, MreB and PBP1 dynamics: 100 µg/mL D-cycloserine, 25 µg/mL fosfomycin, 20 µg/mL penicillin G or 2 µg/mL vancomycin.

### Growth curves

An overnight culture grown in LB medium at 30°C was diluted in fresh LB to OD_600_ 0.01 and incubated at 37°C in tubes to start re-growth. This culture was re-diluted to OD600 0.01 in fresh LB and incubated at 37°C in a plate reader with reading of optical density at 600 nm every 5 min until mid-exponential phase (t0); drugs were added at t0 and incubation was continued in the plate reader.

### Scanning electron microscopy

An overnight culture grown in LB medium at 30°C was diluted in fresh LB to OD600 0.01 and incubated at 37°C until exponential phase. Antibiotics were added and cells allowed to grow for 2 more hours at 37°C before fixation. To this end, a square piece of 1cm² of microscopy slide was placed in the wells of at 24-wells-plate and covered with glutaraldehyde 2% v/v in 0.1 M sodium cacodylate buffer (pH 7,2). 100 µL of cells were added to the wells and incubated 2h at room temperature and then overnight at 4°C. After one wash (15 min) with cacodylate 0,1 M (pH 7,2), dehydration was carried out using ethanol of increasing concentration: 50% (10 min), 70% (10 min), 90% (10 min), 100% (10 min*2). Samples were critical-point dried with liquid CO_2_ as the transition fluid with a Leica CPD300 device and mounted on a 12.8 mm aluminium short pin stub with double-sided sticky and conductive tabs. After desiccation, samples were sputter-coated with an ACE600 device (Leica) in Argon plasma to obtain a 6 nm platinum layer and observed by Scanning Electron Microscopy on a FEG Gemini 500 (Zeiss) microscope at the IBPS platform. Observations were performed in high vacuum (around 1.10^-6^ mbar in the observation chamber), at 3 kV, aperture 20 µm, with high current mode and at around 3.5 mm working distance. Acquisitions were made with the SE (secondary electrons) in lens or SE2 in chamber detectors, driven by SmartSEM software 6.3. Images were recorded with a 1024 x 768 pixels definition and with a pixel dwell time of 1.6 µs and a line averaging of 10.

Antibiotics were used at the following concentrations: fosfomycin 25µg/mL, bacitracin 400µg/mL, D-cycloserine 20 and 100µg/mL, vancomycin 2µg/mL, penicillin G 5µg/mL and 20µg/mL.

### Transmission Electron Microscopy

An overnight culture grown in LB medium at 30°C was diluted in fresh LB to OD_600_ 0.01 and incubated at 37°C until exponential phase. Antibiotics were added and cells grown for 2h at 37°C before collection of the samples by centrifugation (3220 g, 10 min). The samples were processed as previously described (Jacq et al. 2018; Mohamed et al. 2021). Briefly, the pellet (1.4 μl) was dispensed on the 200-μm side of a type A 3-mm gold platelet (Leica Microsystems), covered with the flat side of a type B 3-mm aluminium platelet (Leica Microsystems), and vitrified by high-pressure freezing using an HPM100 system (Leica Microsystems). The samples were then freeze-substituted at −90 °C for 80 h in acetone supplemented with 1% OsO4. The samples were warmed up slowly (1 °C/h) first to −60 °C (AFS2; Leica Microsystems) and after 8 to 12 h the temperature was raised (1 °C/h) to −30 °C. The samples were kept at −30 °C for another 8 to 12 h before being rinsed 4 times in pure acetone. The samples were then infiltrated with gradually increasing concentrations of resin (Embed812, EMS) in acetone (1:2, 1:1, 2:1 [vol/vol]) for 3 h while raising the temperature until it reaches 20 °C. Pure resin was added at room temperature. After polymerization at 60 °C, the samples were cut using an ultramicrotome UC7 (Leica Microsystems) to obtained 80-nm-thin sections. These thin sections were collected on formvar-carbon-coated 100-mesh copper grids, post stained for 5 min with 5% aqueous uranyl acetate, rinsed, and incubated for 2 min with lead citrate. The samples were observed using a CM12 (Philips) or Tecnai G2 Spirit BioTwin (FEI) microscope operating at 120 kV with an Orius SC1000B CCD camera (Gatan).

Cell-wall thickness was measured on x13000 TEM images using Fiji software. At least 3 measures were taken on each cell and averaged; cell-wall thickness was measured on 12 to 25 cells depending on the condition.

Two concentrations were used for each antibiotics: fosfomycin 5 & 25µg/mL, bacitracin 50 & 400µg/mL, D-cycloserine 20 & 100µg/mL, vancomycin 0.25 & 2µg/mL, penicillin G 5 & 20µg/mL chloramphenicol 2µg/mL.

### Timelapses in phase contrast and epifluorescence microscopy

Overnight cultures grown in LB medium at 30°C were diluted in fresh LB to OD_600_ 5.10^-4^ and incubated at 37°C. When cells reached early exponential phase they were stained for 10 min with 10 µg/mL of the membrane dye Nile Red. 1 µL of cells was added to a 1% agarose pad prepared with LB medium containing 10 µg/mL Nile Red into the channel of an Ibidi sticky-slide 0.2 Luer. The mounting was immediately sealed with a coverslip. Cells were allowed to grow for 20 min at 37°C before injection of a drug solution containing 500 µL LB, antibiotic at the appropriate concentration and 10 µg/mL Nile Red. Timelapse acquisitions were realized at 37°C in a temperature-controlled microscope stage by taking images every 2 min in phase contrast and in fluorescence using a 561 nm laser. Microscopy acquisitions were performed using an inverted microscope Nikon Ti-E with a Hamamatsu camera. Antibiotics were used at the following concentrations: fosfomycin 25µg/mL, bacitracin 400µg/mL, D-cycloserine 100µg/mL, vancomycin 2µg/mL, penicillin G 20µg/mL, lysozyme 20µg/mL, chloramphenicol 5µg/mL.

### Morphometric analysis of B. subtilis cells

Fluorescence images of *B. subtilis* cells labelled with Nile Red were semi-automatically segmented to extract the morphology of individual cells. A custom Stardist-based (Schmidt et al. 2018) model was trained to detect the outline of bacterial cells labeled with a membrane marker. Training was

performed using the ZeroCostDL4Mic platform (von Chamier et al. 2021). Predictions were then visually inspected and manually corrected using a custom Napari plugin (https://github.com/aurelien-barbotin/napari-segment-update). To limit manual correction time, only one every 15-time frames was analysed. Cells that were out of focus, dead or impossible to separate from their neighbours were left out of the analysis.

The medial axes of segmented cells were extracted to measure cell width and length, as described in (Ducret et al. 2016). The code is available at https://github.com/aurelien-barbotin/bac_morphology.

### Propidium iodide staining

An overnight culture grown in LB medium at 30°C was diluted in fresh LB +/- 20mM MgSO_4_ to OD600

0.01 and incubated at 37°C. 100 µg/mL D-cycloserine added to exponentially-growing cells. After 1 h incubation with antibiotic, 5 µM propidium iodide was added to the culture before immobilisation of the cells on 1% agarose pad and imaging in epifluorescence microscopy using an inverted microscope Nikon Ti-E with a Hamamatsu camera.

### FDAA staining

(Hsu et al. 2017; Kuru et al. 2019) An overnight culture grown in LB medium at 30°C was diluted in fresh LB to OD_600_ 0.01 and incubated at 37°C. Antibiotics were added to 100 µL aliquots of exponentially-growing cells. After 40 minutes, 0.1 mM sBADA [sulfonated BODIPY-FL 3-amino-D-alanine] was added and cells were allowed to grow for 10 more minutes. Three washings were performed with 500 µL PBS with the corresponding antibiotics before immobilisation of the cells on 1% agarose pad.

Phase contrast and fluorescence (488 nm laser) acquisitions were performed using an inverted microscope Nikon Ti-E with a Hamamatsu camera. Super-resolution images were acquired on an Elyra 7 AxioObserver microscope (Zeiss) and reconstructed using ZEN software (Black Edition, 2012) with the SIM² algorithm, as previously described (Lablaine et al. 2025).

Fluorescence intensity was measured along the sidewall on pseudo-widefield images (prior to SIM image reconstruction) using Fiji, and normalized by the fluorescence of the background.

### MreB TIRF microscopy

TIRF microscopy was essentially realized as described in (Billaudeau et al. 2017). An overnight culture of RCL421 (*gfp-mreB* strain) grown in LB medium at 30°C was diluted in fresh LB to OD_600_ 0.005 and incubated at 37°C until exponential phase. 1 µL of cells was added to a 1% agarose pad prepared with LB medium into the channel of an Ibidi sticky-slide 0.2 Luer. The mounting was immediately sealed with a coverslip and put into the temperature-controlled microscope stage. A first acquisition was realized (t0) and then every 20 min; antibiotics were injected at t20. GFP-MreB were imaged every 0.5 s over 1 min movies, with 100 ms exposure time. Imaging was performed using an inverted microscope (Nikon Ti-E) with an Apo TIRF x100 oil objective (Nikon, NA 1.49).

MreB patches detection and single-particle tracking were realized using Fiji and Matlab softwares, as previously described (Billaudeau et al. 2017, 2020).

### mNeonGreen-PBP1 microscopy

An overnight culture of MK0287 (P_hyperspank_-mNeonGreen-PBP1) grown in LB medium at 30°C was diluted in fresh LB to OD_600_ 0.005 and incubated at 37°C until exponential phase. Antibiotics (D-cycloserine 100µg/mL, vancomycin 2µg/mL) were added to a 100 µL aliquot of culture. After 1h incubation with the antibiotic, 1 µL of cells was added to a 1% agarose pad prepared with LB and antibiotic (same conditions as in the culture). No inducer was used, the leakage from the P_hyperspank_ being sufficient for particle tracking experiments (Cho et al. 2016).

mNeonGreen-PBP1 was imaged every 0.2 s over 1 min movies, with 4 frames per second. Imaging was performed using an inverted microscope (Zeiss SP1).

Images were pre-processed with a sliding window 1 s averaging. PBP1 detection and single-particle tracking were realized using Trackmate (Fiji).

Map of classified trajectories and diffusion coefficient were extracted from a dense mapping analysis performed as described in (Salomon et al. 2020).

## Acknowledgements

The authors thank all members of the Carballido-Lόpez lab for advice and thoughtful discussions throughout this project. Thanks also to Shailab Shrestha for a very useful discussion at the GRC “Bacterial Cell Surface 2022”. We thank Alexis Canette from the IBPS electron microscopy core facility (Sorbonne université - CNRS), and the platform of the Grenoble Instruct-ERIC center for the TEM microscopy (ISBG; UAR 3518 CNRS-CEA-UGA-EMBL) within the Grenoble Partnership for Structural Biology (PSB), supported by FRISBI (ANR-10-INBS-05-02) and GRAL, financed within the University Grenoble Alpes graduate school CBH-EUR-GS (ANR-17-EURE-0003). This work was supported by funding from the European Research Council (ERC) under the European Union’s Horizon 2020 research and innovation programme (ERC Consolidator grant (CoG) BActin 772178 to R.C.L.).

## Authors contributions

R.C.-L. conceptualized, supervised and administrated the project; C.C. and R.C.-L. designed the study, analysed and interpreted data; C.C. conceived experiments and validated results and experiments; C.C., C.-J.R., A.L. and E.B. performed experiments or data collection; L.D. carried out the morphometric analysis; A.B. developed the morphometric analysis; C.B. developed the analysis of MreB and PBP1 dynamics; C.M. contributed to images acquisition by TEM; C.C. and R.C.-L. wrote, revised and edited the manuscript; R.C.-L. administrated and acquired the financial support for the project leading to this publication. All authors read and approved the manuscript.

## Supporting Information

Supplementary Figures S1 to S11

Supplementary Movies S1 to S17

## References

Aghamali, Mina, Mansour Sedighi, Abed Zahedi bialvaei, et al. 2019. ‘Fosfomycin: Mechanisms and the Increasing Prevalence of Resistance’. Journal of Medical Microbiology 68 (1): 11–25. 10.1099/jmm.0.000874.

Atrih, Abdelmadjid, Gerold Bacher, Günter Allmaier, Michael P. Williamson, and Simon J. Foster. 1999. ‘Analysis of Peptidoglycan Structure from Vegetative Cells of *Bacillus Subtilis* 168 and Role of PBP 5 in Peptidoglycan Maturation’. Journal of Bacteriology 181 (13): 3956–66. 10.1128/JB.181.13.3956-3966.1999.

Banzhaf, Manuel, Bart van den Berg van Saparoea, Mohammed Terrak, et al. 2012. ‘Cooperativity of Peptidoglycan Synthases Active in Bacterial Cell Elongation’. Molecular Microbiology 85 (1): 179–94. 10.1111/j.1365-2958.2012.08103.x.

Batson, Sarah, Cesira de Chiara, Vita Majce, et al. 2017. ‘Inhibition of D-Ala:D-Ala Ligase through a Phosphorylated Form of the Antibiotic D-Cycloserine’. Nature Communications 8 (1): 1939. 10.1038/s41467-017-02118-7.

Billaudeau, C., A. Chastanet, and R. Carballido-Lopez. 2020. ‘Processing TIRF Microscopy Images to Characterize the Dynamics and Morphology of Bacterial Actin-like Assemblies’. In CYTOSKELETON DYNAMICS: METHODS AND PROTOCOLS, edited by H. Maiato, vol. 2101. WOS:000664494600010. 10.1007/978-1-0716-0219-5_9.

Billaudeau, C., A. Chastanet, Z. Yao, et al. 2017. ‘Contrasting Mechanisms of Growth in Two Model Rod-Shaped Bacteria’. Nature Communications 8 (June). WOS:000402802900001. 10.1038/ncomms15370.

Billaudeau, C., ZZ Yao, C. Cornilleau, R. Carballido-Lopez, and A. Chastanet. 2019. ‘MreB Forms Subdiffraction Nanofilaments during Active Growth in Bacillus Subtilis’. MBIO 10 (1). WOS:000460314300015. 10.1128/mBio.01879-18.

Bisicchia, Paola, David Noone, Efthimia Lioliou, et al. 2007. ‘The Essential YycFG Two-Component System Controls Cell Wall Metabolism in *Bacillus Subtilis*’. Molecular Microbiology 65 (1): 180– 200. 10.1111/j.1365-2958.2007.05782.x.

Born, Petra, Eefjan Breukink, and Waldemar Vollmer. 2006. ‘In Vitro Synthesis of Cross-Linked Murein and Its Attachment to Sacculi by PBP1A from Escherichia Coli’. The Journal of Biological Chemistry 281 (37): 26985–93. 10.1074/jbc.M604083200.

Brunet, Yannick R., Cameron Habib, Anna P. Brogan, Lior Artzi, and David Z. Rudner. 2022. ‘Intrinsically Disordered Protein Regions Are Required for Cell Wall Homeostasis in Bacillus Subtilis’. *Genes & Development*, ahead of print, October 20. 10.1101/gad.349895.122.

Carballido-López, Rut, Alex Formstone, Ying Li, S. Dusko Ehrlich, Philippe Noirot, and Jeff Errington. 2006. ‘Actin Homolog MreBH Governs Cell Morphogenesis by Localization of the Cell Wall Hydrolase LytE’. Developmental Cell 11 (3): 399–409. 10.1016/j.devcel.2006.07.017.

Chamier, Lucas von, Romain F. Laine, Johanna Jukkala, et al. 2021. ‘Democratising Deep Learning for Microscopy with ZeroCostDL4Mic’. Nature Communications 12 (1): 1. 10.1038/s41467-021-22518-0.

Chastanet, Arnaud, and Rut Carballido-Lopez. 2012. ‘The Actin-like MreB Proteins in Bacillus Subtilis: A New Turn’. Frontiers in Bioscience (Scholar Edition) 4 (June): 1582–606. 10.2741/s354.

Chen, Xin, Suyi Zhong, Yiwei Hou, et al. 2023. ‘Superresolution Structured Illumination Microscopy Reconstruction Algorithms: A Review’. Light: Science & Applications 12 (1): 172. 10.1038/s41377-023-01204-4.

Cho, Hongbaek, Carl N. Wivagg, Mrinal Kapoor, et al. 2016. ‘Bacterial Cell Wall Biogenesis Is Mediated by SEDS and PBP Polymerase Families Functioning Semi-Autonomously’. Nature Microbiology 1 (September): 16172. 10.1038/nmicrobiol.2016.172.

Cleverley, Robert M., Zoe J. Rutter, Jeanine Rismondo, et al. 2019. ‘The Cell Cycle Regulator GpsB Functions as Cytosolic Adaptor for Multiple Cell Wall Enzymes’. Nature Communications 10 (January): 261. 10.1038/s41467-018-08056-2.

Cornilleau, Charlene, Laura Alvarez, Christine Wegler, Cyrille Billaudeau, Felipe Cava, and Rut Carballido-Lopez. 2023. ‘Peptidoglycan Remodeling in Response to Cell Wall Acting Antibiotics in Bacillus Subtilis’. Preprint, bioRxiv, January 23. 10.1101/2023.01.23.525174.

Dajkovic, Alex, Benoit Tesson, Smita Chauhan, et al. 2017. ‘Hydrolysis of Peptidoglycan Is Modulated by Amidation of *Meso* -Diaminopimelic Acid and Mg ^2+^ in *Bacillus Subtilis*: Homeostasis of Peptidoglycan Hydrolysis in *Bacillus Subtilis*’. Molecular Microbiology 104 (6): 972–88. 10.1111/mmi.13673.

Dion, Michael F., Mrinal Kapoor, Yingjie Sun, et al. 2019. ‘Bacillus Subtilis Cell Diameter Is Determined by the Opposing Actions of Two Distinct Cell Wall Synthetic Systems’. Nature Microbiology 4 (8): 1294–305. 10.1038/s41564-019-0439-0.

Domínguez-Cuevas, Patricia, Ida Porcelli, Richard A. Daniel, and Jeff Errington. 2013. ‘Differentiated Roles for MreB-Actin Isologues and Autolytic Enzymes in Bacillus Subtilis Morphogenesis’. Molecular Microbiology 89 (6): 1084–98. 10.1111/mmi.12335.

Dominguez-Escobar, J., A. Chastanet, A. H. Crevenna, V. Fromion, R. Wedlich-Soldner, and R. Carballido-Lopez. 2011. ‘Processive Movement of MreB-Associated Cell Wall Biosynthetic Complexes in Bacteria’. Science 333 (6039): 225–28. 10.1126/science.1203466.

Dörries, Kirsten, Rabea Schlueter, and Michael Lalk. 2014. ‘Impact of Antibiotics with Various Target Sites on the Metabolome of *Staphylococcus Aureus*’. Antimicrobial Agents and Chemotherapy 58 (12): 7151–63. 10.1128/AAC.03104-14.

Ducret, Adrien, Ellen M. Quardokus, and Yves V. Brun. 2016. ‘MicrobeJ, a Tool for High Throughput Bacterial Cell Detection and Quantitative Analysis’. Nature Microbiology 1 (7): 7. 10.1038/nmicrobiol.2016.77.

Emami, Kaveh, Aurelie Guyet, Yoshikazu Kawai, et al. 2017. ‘RodA as the Missing Glycosyltransferase in *Bacillus Subtilis* and Antibiotic Discovery for the Peptidoglycan Polymerase Pathway’. Nature Microbiology 2 (3): 16253. 10.1038/nmicrobiol.2016.253.

Garde, Shambhavi, Pavan Kumar Chodisetti, and Manjula Reddy. 2021. ‘Peptidoglycan: Structure, Synthesis, and Regulation’. EcoSal Plus, ahead of print, January 20. Washington, DC. 10.1128/ecosalplus.ESP-0010-2020.

Garner, E. C., R. Bernard, W. Wang, X. Zhuang, D. Z. Rudner, and T. Mitchison. 2011. ‘Coupled, Circumferential Motions of the Cell Wall Synthesis Machinery and MreB Filaments in *B. Subtilis*’. Science 333 (6039): 222–25. 10.1126/science.1203285.

Halliday, Judy, Declan McKeveney, Craig Muldoon, Premraj Rajaratnam, and Wim Meutermans. 2006. ‘Targeting the Forgotten Transglycosylases’. Biochemical Pharmacology 71 (7): 957–67. 10.1016/j.bcp.2005.10.030.

Hsu, Yen-Pang, Jonathan Rittichier, Erkin Kuru, et al. 2017. ‘Full Color Palette of Fluorescent D -Amino Acids for in Situ Labeling of Bacterial Cell Walls’. Chemical Science 8 (9): 6313–21. 10.1039/C7SC01800B.

Huber, Gerhard, and Georg Nesemann. 1968. ‘Moenomycin, an Inhibitor of Cell Wall Synthesis’. Biochemical and Biophysical Research Communications 30 (1): 7–13. 10.1016/0006-291X(68)90704-3.

Jacq, Maxime, Christopher Arthaud, Sylvie Manuse, et al. 2018. ‘The Cell Wall Hydrolase Pmp23 Is Important for Assembly and Stability of the Division Ring in Streptococcus Pneumoniae’. Scientific Reports 8 (1): 1. 10.1038/s41598-018-25882-y.

Juillot, C. Cornilleau, N. Deboosere, et al. 2021. ‘A High-Content Microscopy Screening Identifies New Genes Involved in Cell Width Control in Bacillus Subtilis’. MSYSTEMS 6 (6). WOS:000790197100008. 10.1128/mSystems.01017-21.

Kahan, F. M., J. S. Kahan, P. J. Cassidy, and H. Kropp. 1974. ‘The Mechanism of Action of Fosfomycin (Phosphonomycin)’. Annals of the New York Academy of Sciences 235 (0): 364–86. 10.1111/j.1749-6632.1974.tb43277.x.

Kawai, Yoshikazu, Kei Asai, and Jeffery Errington. 2009. ‘Partial Functional Redundancy of MreB Isoforms, MreB, Mbl and MreBH, in Cell Morphogenesis of *Bacillus Subtilis*’. Molecular Microbiology 73 (4): 719–31. 10.1111/j.1365-2958.2009.06805.x.

Kawai, Yoshikazu, Richard A. Daniel, and Jeffery Errington. 2009. ‘Regulation of Cell Wall Morphogenesis in *Bacillus Subtilis* by Recruitment of PBP1 to the MreB Helix’. Molecular Microbiology 71 (5): 1131–44. 10.1111/j.1365-2958.2009.06601.x.

Kawai, Yoshikazu, Maki Kawai, Eilidh Sohini Mackenzie, et al. 2023. ‘On the Mechanisms of Lysis Triggered by Perturbations of Bacterial Cell Wall Biosynthesis’. Nature Communications 14 (July): 4123. 10.1038/s41467-023-39723-8.

Kawai, Yoshikazu, Katarzyna Mickiewicz, and Jeff Errington. 2018. ‘Lysozyme Counteracts β-Lactam Antibiotics by Promoting the Emergence of L-Form Bacteria’. Cell 172 (5): 1038–1049.e10. 10.1016/j.cell.2018.01.021.

Kitahara, Yuki, Enno R. Oldewurtel, Sean Wilson, Yingjie Sun, Ethan C. Garner, and Sven van Teeffelen. 2021. Cell-Envelope Synthesis Is Required for Surface-to-Mass Coupling, Which Determines Dry-Mass Density in Bacillus Subtilis. Preprint. Microbiology. 10.1101/2021.05.05.442853.

Kuru, Erkin, H. Velocity Hughes, Pamela J. Brown, et al. 2012. ‘In Situ Probing of Newly Synthesized Peptidoglycan in Live Bacteria with Fluorescent D-Amino Acids’. Angewandte Chemie (International Ed. in English) 51 (50): 12519–23. 10.1002/anie.201206749.

Kuru, Erkin, Atanas Radkov, Xin Meng, et al. 2019. ‘Mechanisms of Incorporation for D -Amino Acid Probes That Target Peptidoglycan Biosynthesis’. ACS Chemical Biology 14 (12): 2745–56. 10.1021/acschembio.9b00664.

Lablaine, Armand, Dimitri Juillot, Ciarán Condon, and Rut Carballido-López. 2025. ‘Real-Time Nanoscale Investigation of Spore Coat Assembly in Bacillus Subtilis’. Communications Biology 8 (1): 1131. 10.1038/s42003-025-08522-w.

Lee, Timothy K., Kevin Meng, Handuo Shi, and Kerwyn Casey Huang. 2016. ‘Single-Molecule Imaging Reveals Modulation of Cell Wall Synthesis Dynamics in Live Bacterial Cells’. Nature Communications 7 (October): 13170. 10.1038/ncomms13170.

Mainardi, Jean-Luc, Régis Villet, Timothy D. Bugg, Claudine Mayer, and Michel Arthur. 2008. ‘Evolution of Peptidoglycan Biosynthesis under the Selective Pressure of Antibiotics in Gram-Positive Bacteria’. FEMS Microbiology Reviews 32 (2): 386–408. 10.1111/j.1574-6976.2007.00097.x.

Mandelstam, J., and H. J. Rogers. 1959. ‘The Incorporation of Amino Acids into the Cell-Wall Mucopeptide of Staphylococci and the Effect of Antibiotics on the Process’. Biochemical Journal 72 (4): 654–62. 10.1042/bj0720654.

McPherson, Derrell C., and David L. Popham. 2003. ‘Peptidoglycan Synthesis in the Absence of Class A Penicillin-Binding Proteins in *Bacillus Subtilis*’. Journal of Bacteriology 185 (4): 1423–31. 10.1128/JB.185.4.1423-1431.2003.

Meeske, Alexander J., Eammon P. Riley, William P. Robins, et al. 2016. ‘SEDS Proteins Are a Widespread Family of Bacterial Cell Wall Polymerases’. Nature 537 (7622): 634–38. 10.1038/nature19331.

Meisner, Jeffrey, Paula Montero Llopis, Lok-To Sham, Ethan Garner, Thomas G. Bernhardt, and David Z. Rudner. 2013. ‘FtsEX Is Required for CwlO Peptidoglycan Hydrolase Activity during Cell Wall Elongation in *Bacillus Subtilis*: FtsEX Governs CwlO Activity’. Molecular Microbiology 89 (6): 1069–83. 10.1111/mmi.12330.

Mohamed, Ahmed M. T., Helena Chan, Johana Luhur, et al. 2021. ‘Chromosome Segregation and Peptidoglycan Remodeling Are Coordinated at a Highly Stabilized Septal Pore to Maintain Bacterial Spore Development’. Developmental Cell 56 (1): 36–51.e5. 10.1016/j.devcel.2020.12.006.

Murphy, Shannon G., Andrew N. Murtha, Ziyi Zhao, et al. 2021. ‘Class A Penicillin-Binding Protein-Mediated Cell Wall Synthesis Promotes Structural Integrity during Peptidoglycan Endopeptidase Insufficiency in Vibrio Cholerae’. mBio, ahead of print, April 6. 1752 N St., N.W., Washington, DC. 10.1128/mBio.03596-20.

Murray, Thomas, David L. Popham, and Peter Setlow. 1998. ‘Bacillus Subtilis Cells Lacking Penicillin-Binding Protein 1 Require Increased Levels of Divalent Cations for Growth’. Journal of Bacteriology 180 (17): 4555–63. https://www.ncbi.nlm.nih.gov/pmc/articles/PMC107467/.

Ostash, Bohdan, and Suzanne Walker. 2010. ‘Moenomycin Family Antibiotics: Chemical Synthesis, Biosynthesis, and Biological Activity’. Natural Product Reports 27 (11): 1594. 10.1039/c001461n.

Patel, Yesha, Heng Zhao, and John D. Helmann. 2020. ‘A Regulatory Pathway That Selectively Up-Regulates Elongasome Function in the Absence of Class A PBPs’. eLife 9 (September): e57902. 10.7554/eLife.57902.

Pazos, Manuel, and Katharina Peters. 2019. ‘Peptidoglycan’. In *Bacterial Cell Walls and Membranes*, edited by Andreas Kuhn, vol. 92. Subcellular Biochemistry. Springer International Publishing. 10.1007/978-3-030-18768-2_5.

Pedro, Miguel A. de, and Felipe Cava. 2015. ‘Structural Constraints and Dynamics of Bacterial Cell Wall Architecture’. Frontiers in Microbiology 6 (May). 10.3389/fmicb.2015.00449.

Popham, D. L., and P. Setlow. 1995. ‘Cloning, Nucleotide Sequence, and Mutagenesis of the Bacillus Subtilis ponA Operon, Which Codes for Penicillin-Binding Protein (PBP) 1 and a PBP-Related Factor’. Journal of Bacteriology 177 (2): 326–35. 10.1128/jb.177.2.326-335.1995.

Rueff, AS, A. Chastanet, J. Dominguez-Escobar, et al. 2014. ‘An Early Cytoplasmic Step of Peptidoglycan Synthesis Is Associated to MreB in Bacillus Subtilis’. Molecular Microbiology 91 (2): 348–62. WOS:000329449600010. 10.1111/mmi.12467.

Salomon, Antoine, Cesar Augusto Valades-Cruz, Ludovic Leconte, and Charles Kervrann. 2020. ‘Dense Mapping of Intracellular Diffusion and Drift from Single-Particle Tracking Data Analysis’. ICASSP 2020 - 2020 IEEE International Conference on Acoustics, Speech and Signal Processing (ICASSP), May, 966–70. 10.1109/ICASSP40776.2020.9054576.

Sarkar, Paramita, Venkateswarlu Yarlagadda, Chandradhish Ghosh, and Jayanta Haldar. 2017. ‘A Review on Cell Wall Synthesis Inhibitors with an Emphasis on Glycopeptide Antibiotics’. MedChemComm 8 (3): 516–33. 10.1039/c6md00585c.

Sassine, Jad, Joana Sousa, Michael Lalk, Richard A. Daniel, and Waldemar Vollmer. 2020. ‘Cell Morphology Maintenance in *Bacillus Subtilis* through Balanced Peptidoglycan Synthesis and Hydrolysis’. Scientific Reports 10 (1): 17910. 10.1038/s41598-020-74609-5.

Sassine, Jad, Meizhu Xu, Karzan R. Sidiq, Robyn Emmins, Jeff Errington, and Richard A. Daniel. 2017. ‘Functional Redundancy of Division Specific Penicillin-binding Proteins in Bacillus Subtilis’. Molecular Microbiology 106 (2): 304–18. 10.1111/mmi.13765.

Sauvage, Eric, Frédéric Kerff, Mohammed Terrak, Juan A. Ayala, and Paulette Charlier. 2008. ‘The Penicillin-Binding Proteins: Structure and Role in Peptidoglycan Biosynthesis’. FEMS Microbiology Reviews 32 (2): 234–58. 10.1111/j.1574-6976.2008.00105.x.

Scheffers, Dirk-Jan, and Mariana G. Pinho. 2005. ‘Bacterial Cell Wall Synthesis: New Insights from Localization Studies’. Microbiology and Molecular Biology Reviews: MMBR 69 (4): 585–607. 10.1128/MMBR.69.4.585-607.2005.

Schirner, Kathrin, Ye-Jin Eun, Mike Dion, et al. 2015. ‘Lipid-Linked Cell Wall Precursors Regulate Membrane Association of Bacterial Actin MreB’. Nature Chemical Biology 11 (1): 38–45. 10.1038/nchembio.1689.

Schmidt, Uwe, Martin Weigert, Coleman Broaddus, and Gene Myers. 2018. ‘Cell Detection with Star-Convex Polygons’. In Medical Image Computing and Computer Assisted Intervention – MICCAI 2018, edited by Alejandro F. Frangi, Julia A. Schnabel, Christos Davatzikos, Carlos Alberola-López, and Gabor Fichtinger. Lecture Notes in Computer Science. Springer International Publishing. 10.1007/978-3-030-00934-2_30.

Schneider, Tanja, and Hans-Georg Sahl. 2010. ‘An Oldie but a Goodie - Cell Wall Biosynthesis as Antibiotic Target Pathway’. International Journal of Medical Microbiology: IJMM 300 (2–3): 161–69. 10.1016/j.ijmm.2009.10.005.

Sharifzadeh, Shabnam, Felix Dempwolff, Daniel B. Kearns, and Erin E. Carlson. 2020. ‘Harnessing β-Lactam Antibiotics for Illumination of the Activity of Penicillin-Binding Proteins in Bacillus Subtilis’. ACS Chemical Biology 15 (5): 1242–51. 10.1021/acschembio.9b00977.

Smith, Thomas J., Steve A. Blackman, and Simon J. Foster. 2000. ‘Autolysins of Bacillus Subtilis: Multiple Enzymes with Multiple Functions’. *Microbiology (Reading*, England) 146 (Pt 2) (February): 249–62. 10.1099/00221287-146-2-249.

Strange, R. E., and L. H. Kent. 1959. ‘The Isolation, Characterization and Chemical Synthesis of Muramic Acid’. The Biochemical Journal 71 (2): 333–39. 10.1042/bj0710333.

Straume, Daniel, Katarzyna Wiaroslawa Piechowiak, Silje Olsen, et al. 2020. ‘Class A PBPs Have a Distinct and Unique Role in the Construction of the Pneumococcal Cell Wall’. Proceedings of the National Academy of Sciences 117 (11): 6129–38. 10.1073/pnas.1917820117.

Sun, Yingjie, Sylvia Hürlimann, and Ethan Garner. 2023. ‘Growth Rate Is Modulated by Monitoring Cell Wall Precursors in Bacillus Subtilis’. Nature Microbiology 8 (3): 469–80. 10.1038/s41564-023-01329-7.

Teeffelen, S. van, S. Wang, L. Furchtgott, et al. 2011. ‘The Bacterial Actin MreB Rotates, and Rotation Depends on Cell-Wall Assembly’. Proceedings of the National Academy of Sciences 108 (38): 15822–27. 10.1073/pnas.1108999108.

Terrak, Mohammed, Tushar K. Ghosh, Jean Van Heijenoort, et al. 1999. ‘The Catalytic, Glycosyl Transferase and Acyl Transferase Modules of the Cell Wall Peptidoglycan-Polymerizing Penicillin-Binding Protein 1b of Escherichia Coli’. Molecular Microbiology 34 (2): 350–64. 10.1046/j.1365-2958.1999.01612.x.

Tesson, Benoit, Alex Dajkovic, Ruth Keary, Christian Marlière, Christine C. Dupont-Gillain, and Rut Carballido-López. 2022. ‘Magnesium Rescues the Morphology of Bacillus Subtilis mreB Mutants through Its Inhibitory Effect on Peptidoglycan Hydrolases’. Scientific Reports 12 (1): 1. 10.1038/s41598-021-04294-5.

Tomasz, A. 1979. ‘The Mechanism of the Irreversible Antimicrobial Effects of Penicillins: How the Beta-Lactam Antibiotics Kill and Lyse Bacteria’. Annual Review of Microbiology 33: 113–37. 10.1146/annurev.mi.33.100179.000553.

Van Heijenoort, Y., M. Derrien, and J. Van Heijenoort. 1978. ‘Polymerization by Transglycosylation in the Biosynthesis of the Peptidoglycan of Escherichia Coli K 12 and Its Inhibition by Antibiotics’. FEBS Letters 89 (1): 141–44. 10.1016/0014-5793(78)80540-7.

Vigouroux, Antoine, Baptiste Cordier, Andrey Aristov, et al. 2020. ‘Class-A Penicillin Binding Proteins Do Not Contribute to Cell Shape but Repair Cell-Wall Defects’. eLife 9 (January): e51998. 10.7554/eLife.51998.

Vollmer, Waldemar, Didier Blanot, and Miguel A. De Pedro. 2008. ‘Peptidoglycan Structure and Architecture’. FEMS Microbiology Reviews 32 (2): 149–67. 10.1111/j.1574-6976.2007.00094.x.

Vollmer, Waldemar, Bernard Joris, Paulette Charlier, and Simon Foster. 2008. ‘Bacterial Peptidoglycan (Murein) Hydrolases’. FEMS Microbiology Reviews 32 (2): 259–86. 10.1111/j.1574-6976.2007.00099.x.

Walsh, C. T. 1989. ‘Enzymes in the D-Alanine Branch of Bacterial Cell Wall Peptidoglycan Assembly’. Journal of Biological Chemistry 264 (5): 2393–96. 10.1016/S0021-9258(19)81624-1.

Wilson, Sean, and Ethan Garner. 2021. An Exhaustive Multiple Knockout Approach to Understanding Cell Wall Hydrolase Function in Bacillus Subtilis. 10.1101/2021.02.18.431929.

Zhao, Heng, Vaidehi Patel, John D. Helmann, and Tobias Dörr. 2017. ‘Don’t Let Sleeping Dogmas Lie: New Views of Peptidoglycan Synthesis and Its Regulation: New Views on Peptidoglycan Synthesis’. Molecular Microbiology 106 (6): 847–60. 10.1111/mmi.13853.

