## Supplementary Figures for "Differential impact of cell wall antibiotics on the Rod complex and aPBPs in *Bacillus subtilis:* Insights into the peptidoglycan elongation machineries"

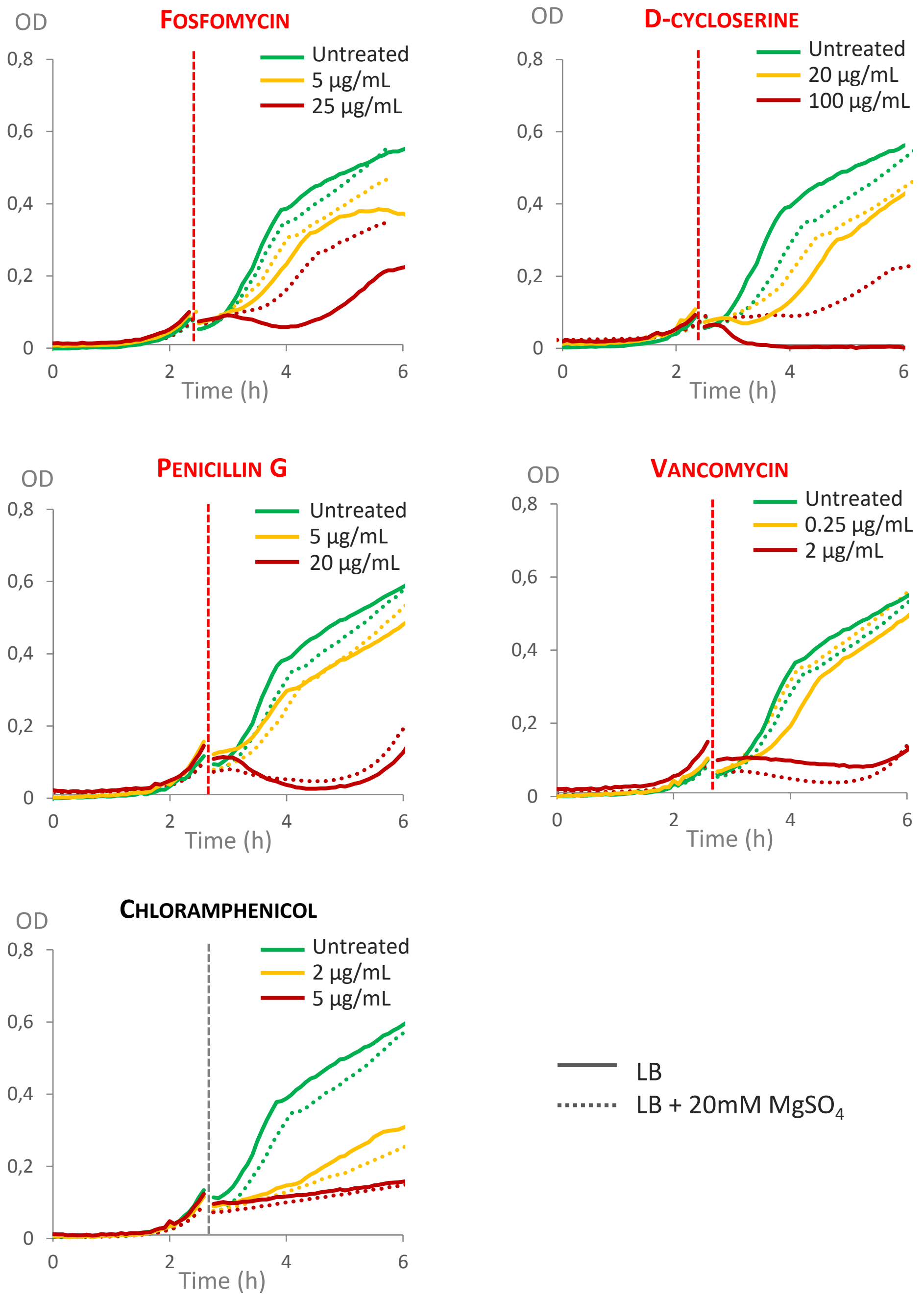


**Fig. S1 - Growth curves with addition of antibiotic in exponential phase**

Growth curves of the wild-type strain 168 growing at 37°C in LB medium (plain lines) and LB + 20 mM $MgSO_4$ (dotted lines) in the presence of fosfomycin, D-cycloserine, penicillin G, vancomycin and choramphenicol. The antibiotics were added at the indicated concentrations when cultures reached exponential phase (dashed line).

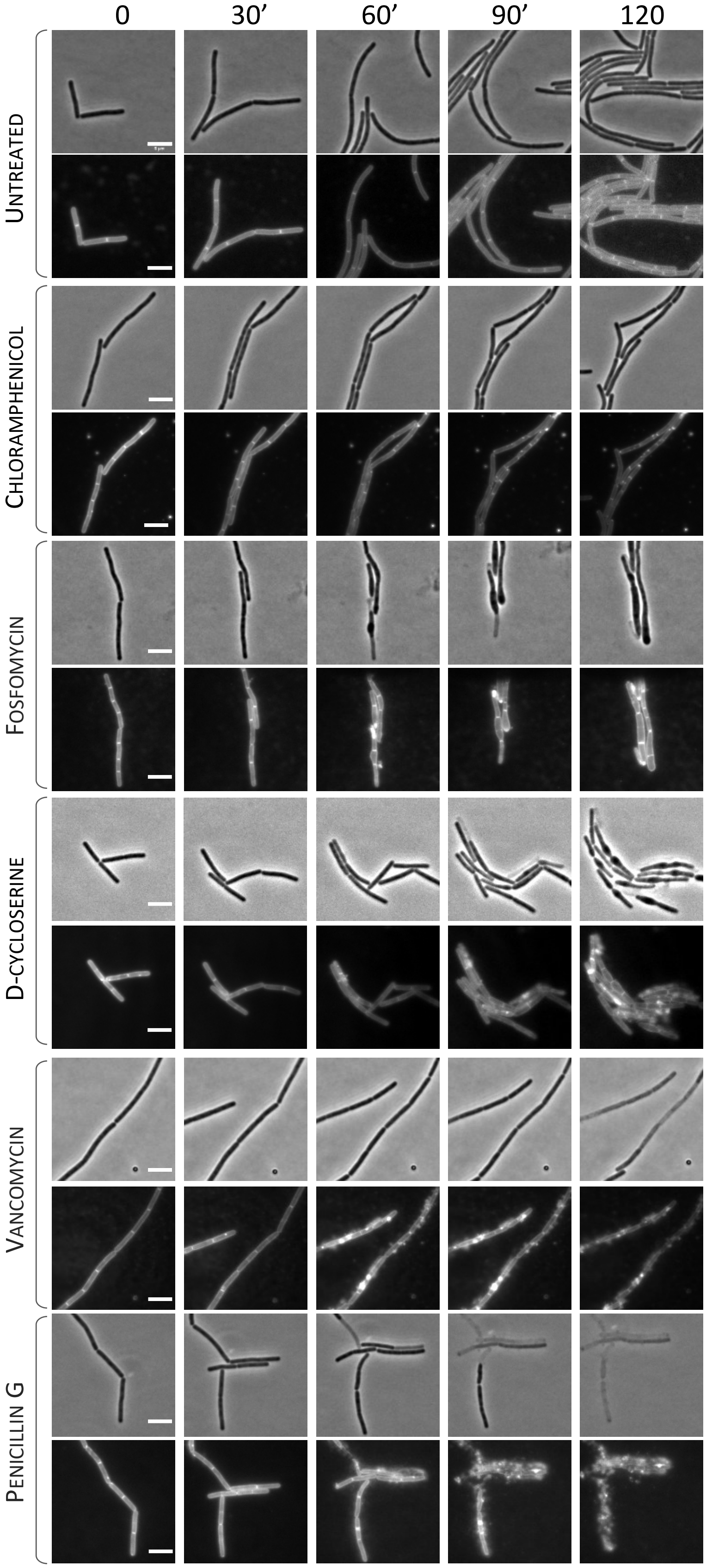


**Fig. S2. Effect of the antibiotics on cell growth and morphology**

Cells were grown in LB medium at 37°C until exponential phase, stained with 10 µg/mL NileRed as membrane dye and spotted on agarose pads prepared in LB with 10 µg/mL NileRed. Cells were allowed to adapt for 20 min and antibiotics were added at the following concentrations: fosfomycin 25 µg/mL, D-cycloserine 100 µg/mL, vancomycin 2 µg/mL, penicillin G 20 µg/mL, chloramphenicol 5 µg/mL. Phase contrast (*Top panels*) and epifluorescence images (*Bottom panels*) of membrane-stained cells were acquired every 2 min for 2 hours (see corresponding Suppl. movies S1-S6). One image acquired every 30 min is shown here. Scale bars, 5 µm.

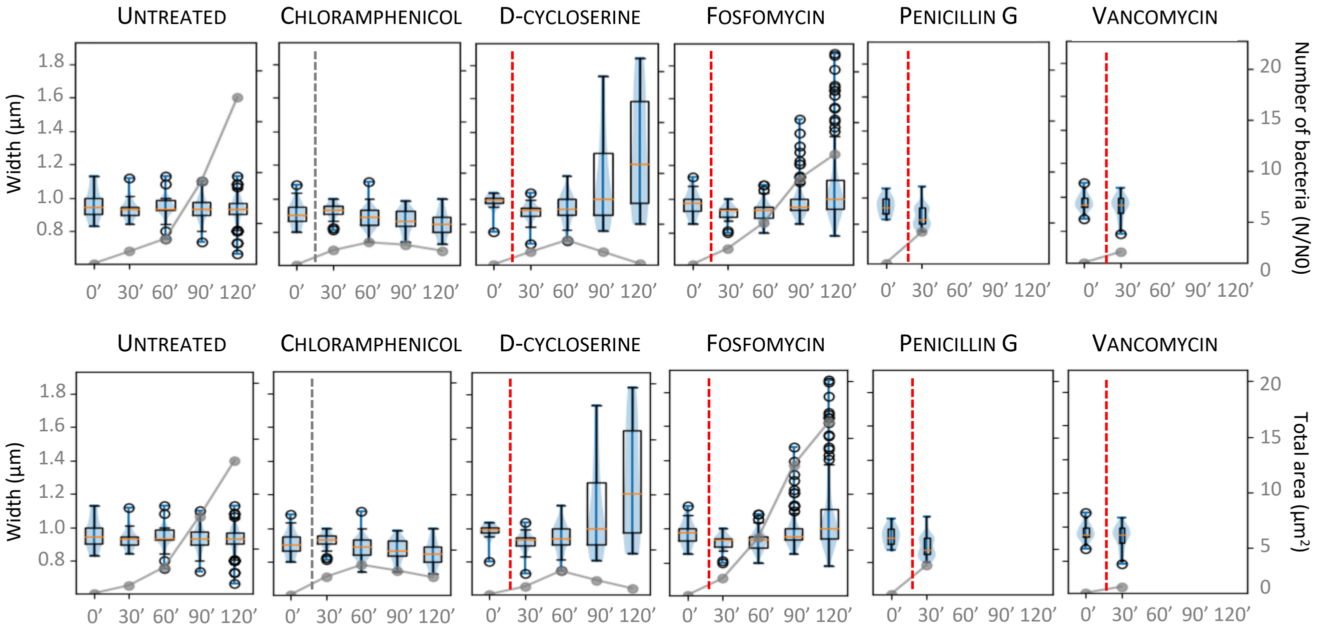


**Fig. S3 – Analysis of cells growth and morphology during antibiotic treatment on agarose pads.**

Cells were grown in LB medium at 37°C until exponential phase, stained with 10 µg/mL NileRed as membrane dye and spotted on agarose pads prepared in LB with 10 µg/mL NileRed. Cells were allowed to settle in the pad for 20 min and antibiotics were added (dashed line) at the following concentrations: fosfomycin 25 µg/mL, D-cycloserine 100 µg/mL, vancomycin 2 µg/mL, penicillin G 20 µg/mL, chloramphenicol 5 µg/mL. The cell diameter (*Left axis*) and the number of cells (*Top panels, right axis*) or total colony area (*Bottom panels, right axis*) were quantified from time-lapses microscopy movies (see **Fig. S2** and **Suppl. movies S1-S6**) and are displayed as boxplots and graphs, respectively. The number of cells was normalized by the number of cells at t0 (N/N0).

Cornilleau *et al* **Fig. S4**

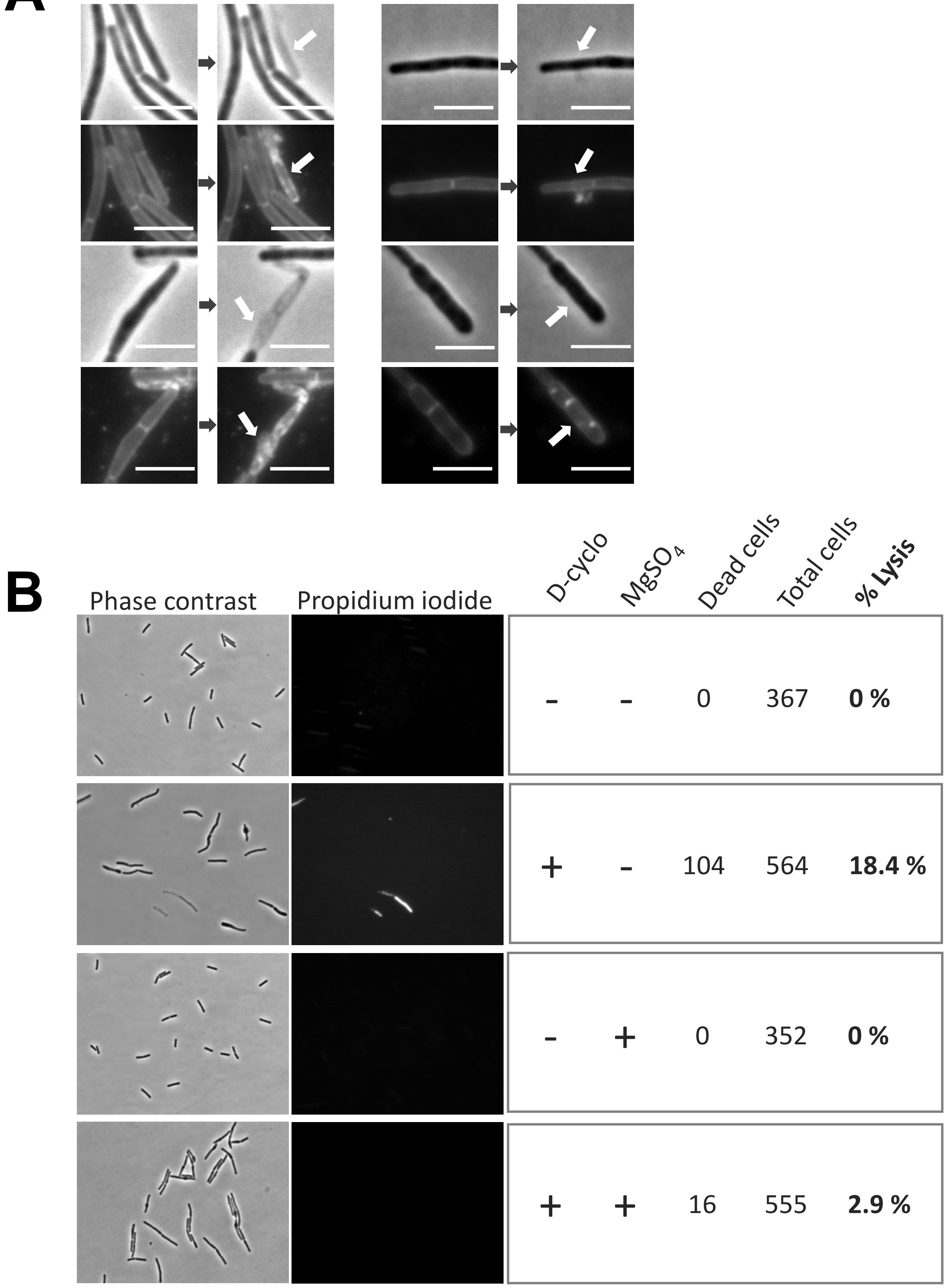


**Fig. S4. Bulging of D-cycloserine and fosfomycin-treated cells is independent of the elevated lysis phenotype.**

**A.** Cells were grown in LB medium at 37°C until exponential phase, stained with 10 µg/mL NileRed as membrane dye and spotted on agarose pads prepared in LB with 10 µg/mL NileRed. Cells were allowed to adapt for 20 min before addition of antibiotic: fosfomycin (25 µg/mL) or D-cycloserine (100 µg/mL). Phase contrast and epifluorescence (membrane staining) images were taken every 2 min during 2 h. White arrows indicated cells that lysed during the 2 minutes interval between acquisitions. Scale bar: 5 µm.

**B**. Phase contrast and propidium iodide labelling of the wt strain, grown in LB +/- 20 mM $MgSO_4$, untreated and treated with 100 µg/mL D-cycloserine during 1 h (addition of antibiotic in exponential phase of growth). The same settings are used for all fluorescent images. The total cells and the dead cells were comptabilized based on phase contrast and propidium iodide labelling to calculate the percentage of lysis: % lysis = 100*((dead cells)/(total cells)).

Cornilleau *et al* **Fig S5**

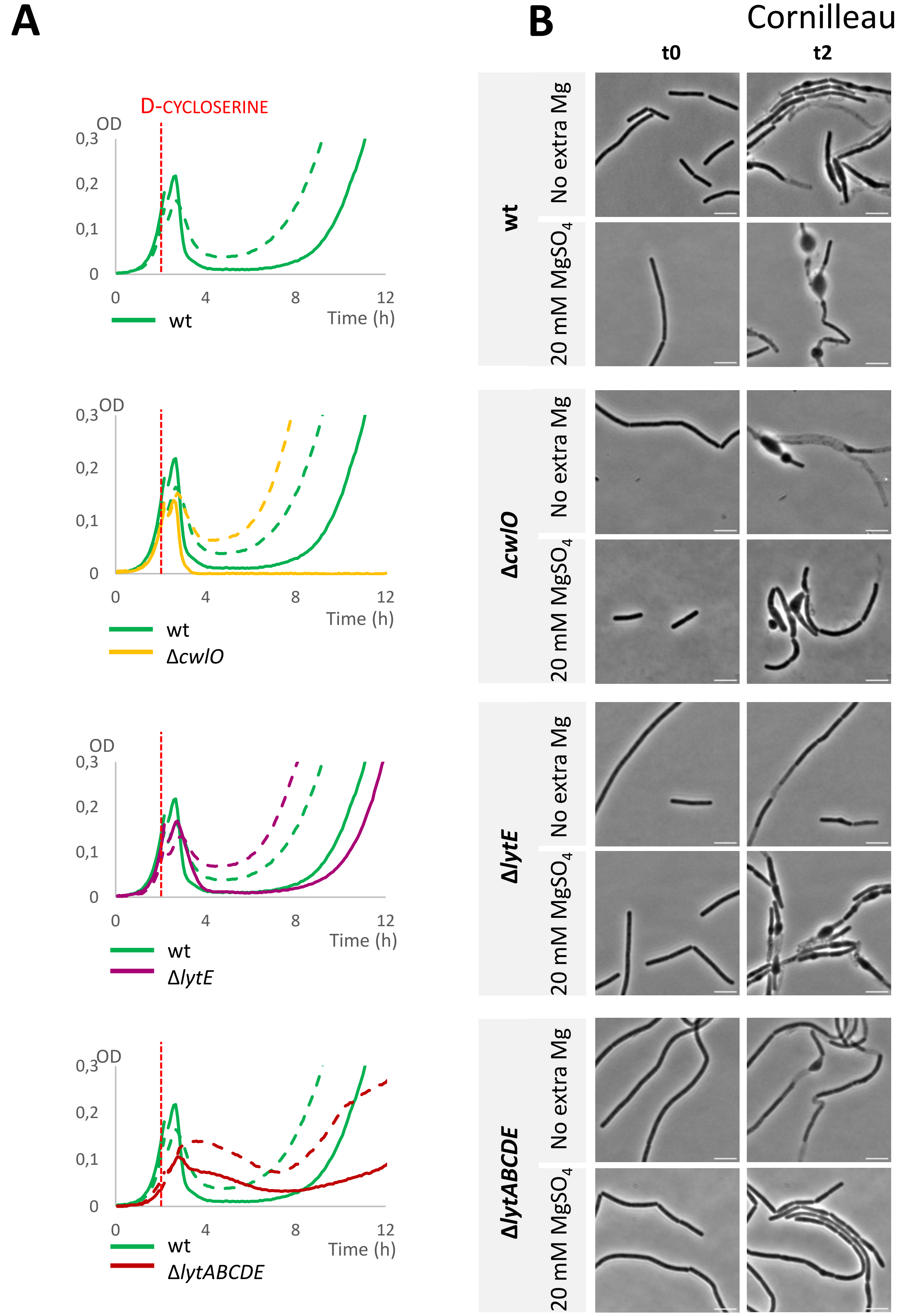


**Fig. S5 - Antibiotic-induced cell bulging is partially dependent (?) on deregulated PG hydrolytic activity**

**A.** Growth curves of the wild-type strain 168, Δ*cwlO*, Δ*lytE* and Δ*lytABCDE* mutants growing at 37°C in LB medium (plain lines) and LB + 20 mM $MgSO_4$ (dotted lines) in the presence of 100 µg/mL D-cycloserine, added when cultures reached exponential phase (red dashed line).

**B.** Phase contrast image of the wild-type strain 168, Δ*cwlO*, Δ*lytE* and Δ*lytABCDE* mutants, grown in LB +/- 20 mM $MgSO_4$, before (t0) and after 2 h (t2) exposure to 25 µg/mL D-cycloserine. Scale bar: 5 µm. See corresponding **Suppl. movies S7-S13**.

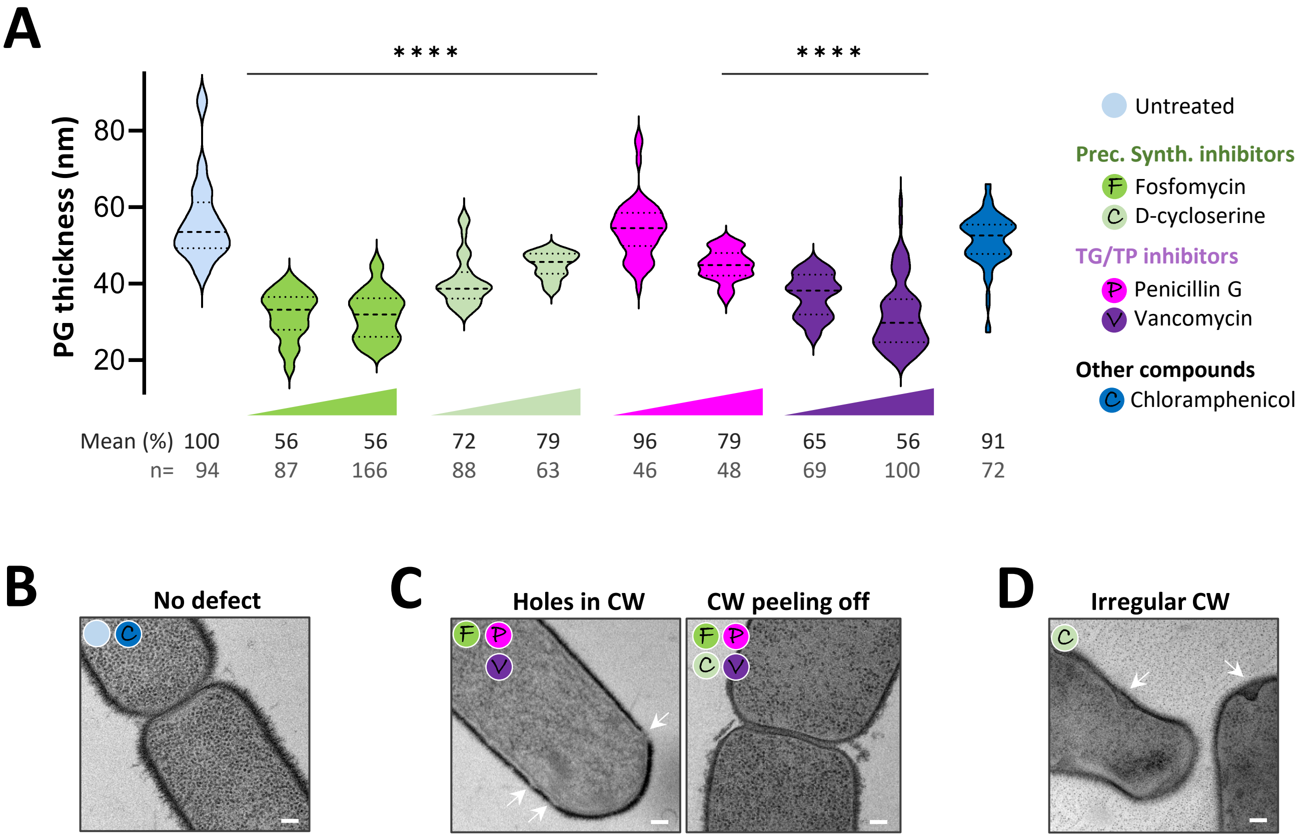


**Fig. S6 - Cell bulging is suspected to be caused by an irregular cell wall**

**A.** Cell wall thickness is impacted by antibiotic treatment. Cell wall thickness (in nm) measured on TEM images and mean thickness expressed as % of the control. Two concentrations were used for each antibiotic: fosfomycin 5 & 25µg/mL, D-cycloserine 20 & 100µg/mL, vancomycin 0.25 & 2µg/mL, penicillin G 5 & 20µg/mL, chloramphenicol 2µg/mL. Significance was evaluated by t-test and indicated as "*".

**B, C, D.** Characteristic structures visibles in TEM: TEM images at x13000 magnification (scale bar: 100 nm), after 2 h incubation with antibiotics. Arrows are pointing at cell wall defects. Antibiotics associated with each phenotype are symbolised with coloured circles.

Cornilleau *et al.* **Fig. S7**

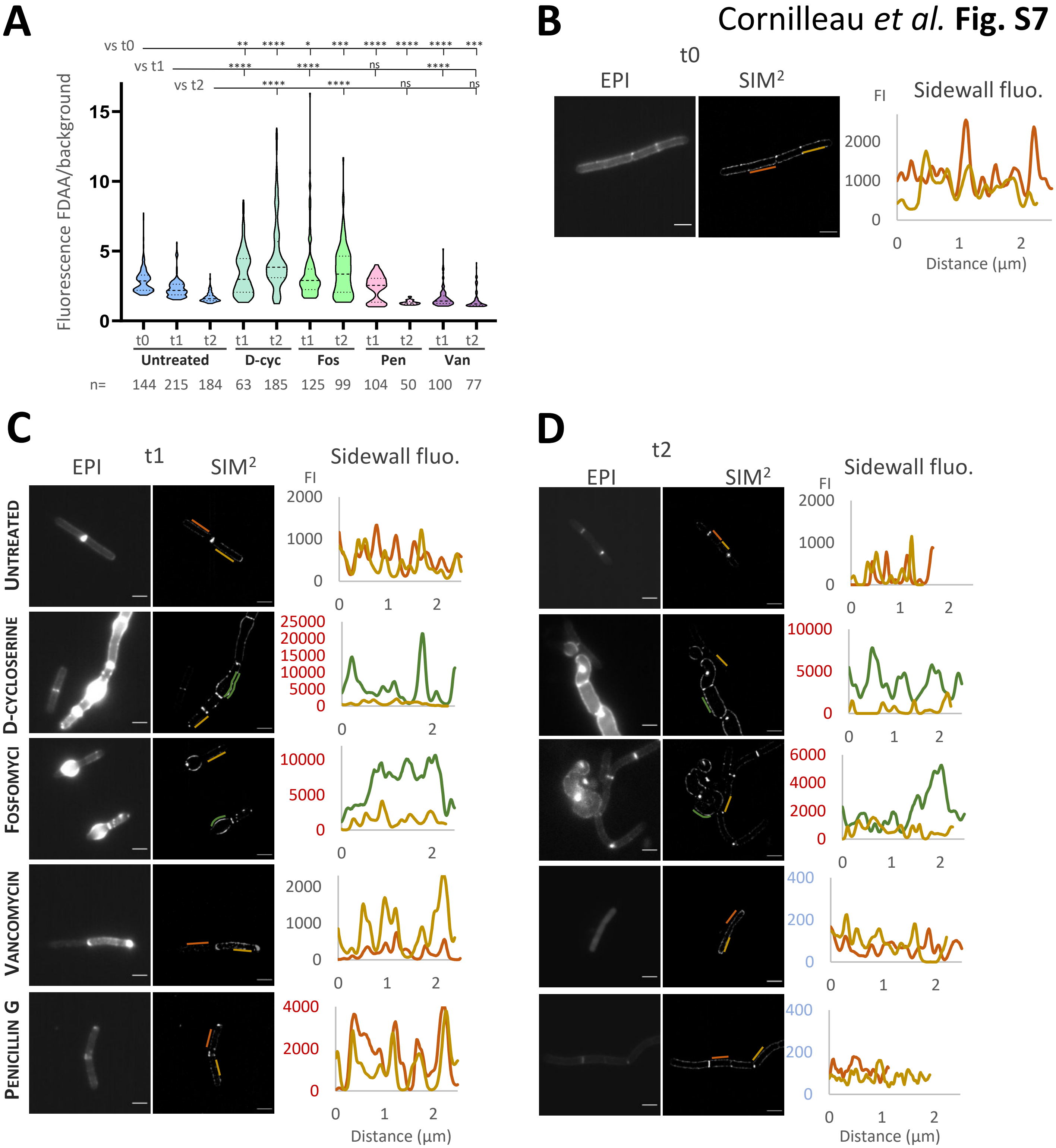


**Fig. S7 - FDAA labelling of *B. subtilis* revealed active PG synthesis in cells treated by inhibitors of PG precursors synthesis**

Antibiotics were used at the following concentrations: 100 µg/mL D-cycloserine (D-cyc), 25 µg/mL fosfomycin (Fos), 20 µg/mL penicillin G (Pen) or 2 µg/mL vancomycin (Van).

**A.** Quantification of sBADA fluorescence intensity measured along the sidewall and normalized by the background. Cells were grown until exponential before addition of antibiotics (t0). sBADA incorporation was assessed at t0 and after 1h (t1) and 2h (t2) of antibiotic treatment. Significance in represented as “*” for each condition against the untreated control at t0, t1 and t2.

**B, C, D.** Epifluorescence and SIM² images of sBADA labelling of *B. subtilis* wt cells before and after addition of antibiotic (scale bar: 2 µm). The intensity profiles (fluorescence intensity FI along the sidewall) are shown for two cells of each condition. **B.** Before addition of antibiotic (t0). **C.** At 1 h (t1) after addition of antibiotic. **D.** At 2 h (t2) after addition of antibiotic.

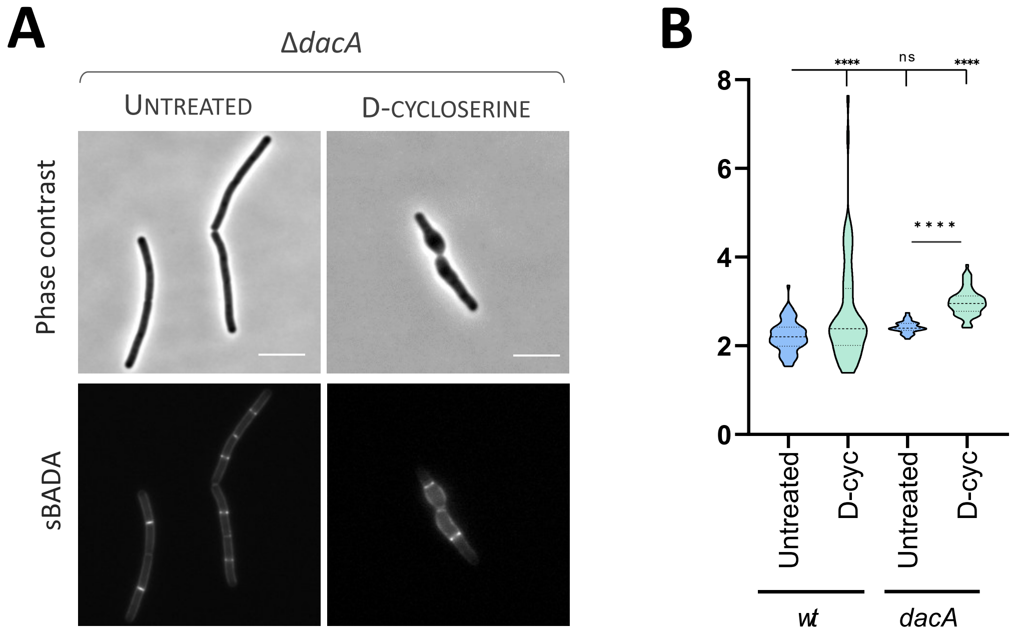


**Fig. S8 - FDAA labelling of *dacA* deletion mutant**

Cell were grown until exponential phase before addition of 100 µg/mL D-cycloserine (D-cyc) and further incubation during 1 h.

**A.** Phase contrast and sBADA labelling of the *dacA* mutant. Same settings for all fluorescent images (scale bar: 5 µm).

**B.** Quantification of sBADA fluorescence intensity measured along the sidewall and normalized by the background, for the wt cells and *dacA* mutant. Significance in represented as "*".

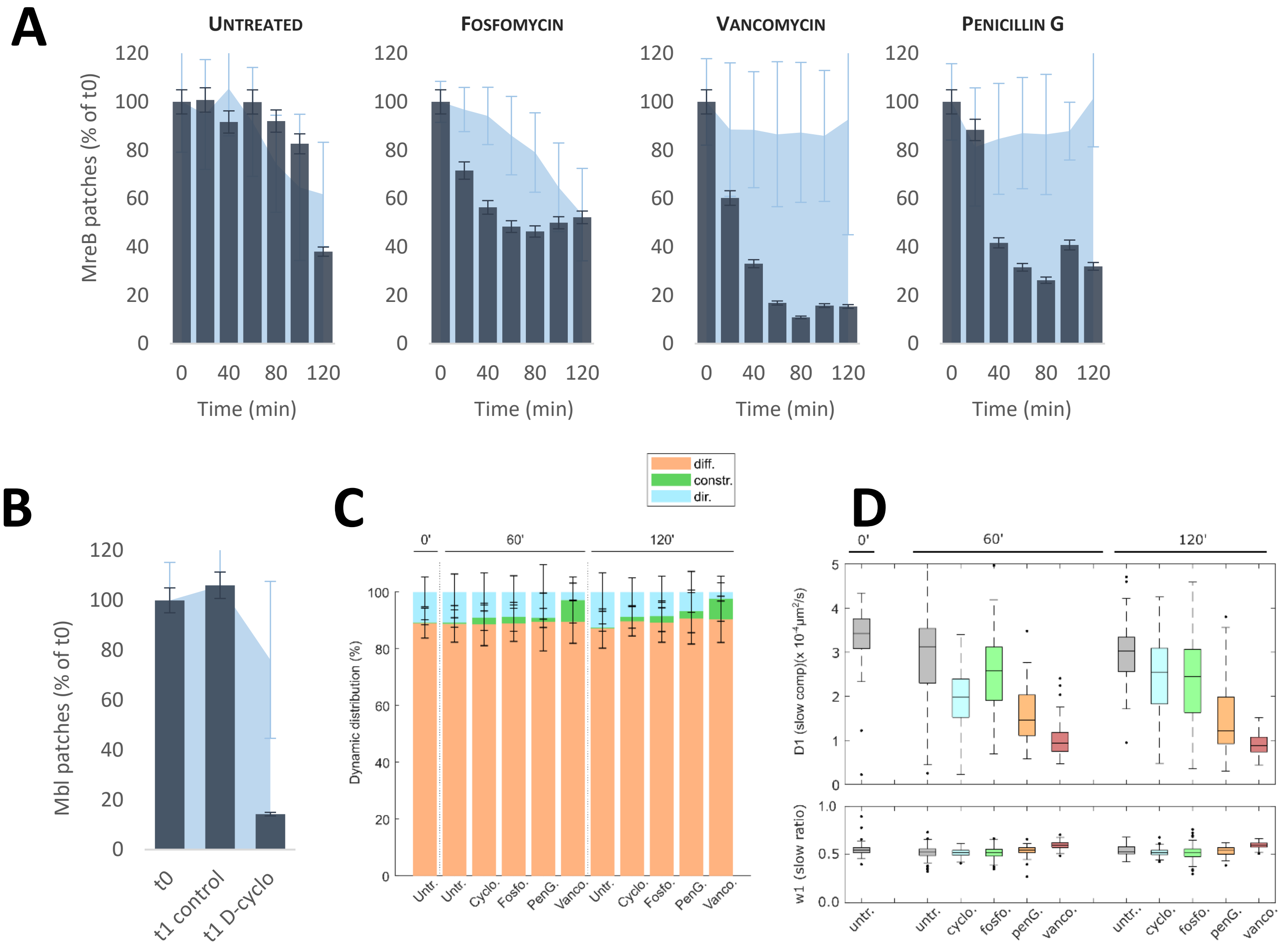


**Fig. S9 - MreB and Mbl are affected by CW antibiotics**

**A.** Dynamic behaviour and density of MreB patches for an untreated control and for cells treated with 25 µg/mL fosfomycin, 2 µg/mL vancomycin or 20 µg/mL penicillin G. The dynamic behaviour is based on MSD analysis, only the percentage of directed patches (normalized by the percentage at t0) is represented here (dark blue histograms). The density of MreB patches is represented as a percentage of the density at t0 (light blue area).

**B.** Dynamic behaviour and density of Mbl patches for an untreated control and for cells treated with 100 µg/mL D-cycloserine. Cells were grown until exponential phase (t0) before addition of D-cycloserine. GFP-Mbl was imaged in TIRF microscopy at t0 and 1h (t1) after. The dynamic behaviour of Mbl patches is based on MSD analysis, only the percentage of directed patches (normalized by the percentage at t0) is represented here (dark blue histograms). The density of Mbl patches is represented as a percentage of the density at t0 (light blue area).

**C.** Dynamic distribution of PBP1 for an untreated control and for cells treated with 25 µg/mL fosfomycin, 100 µg/mL D-cycloserine, 2 µg/mL vancomycin or 20 µg/mL penicillin G.

**D.** Diffusion was quantified at single cell levels (or filament levels if impossible to resolve) using CDF analysis with two components D1 and D2, resp. slow and fast components, and w1 is the weight of slow component in the population.

**A**

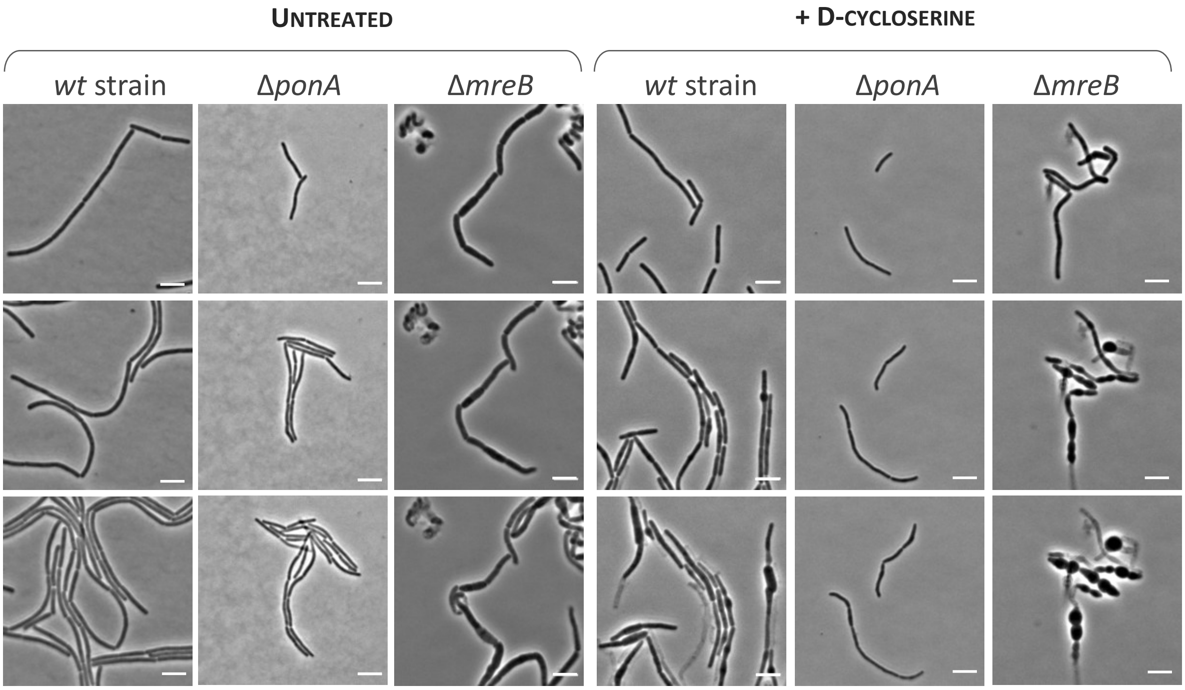


**B**

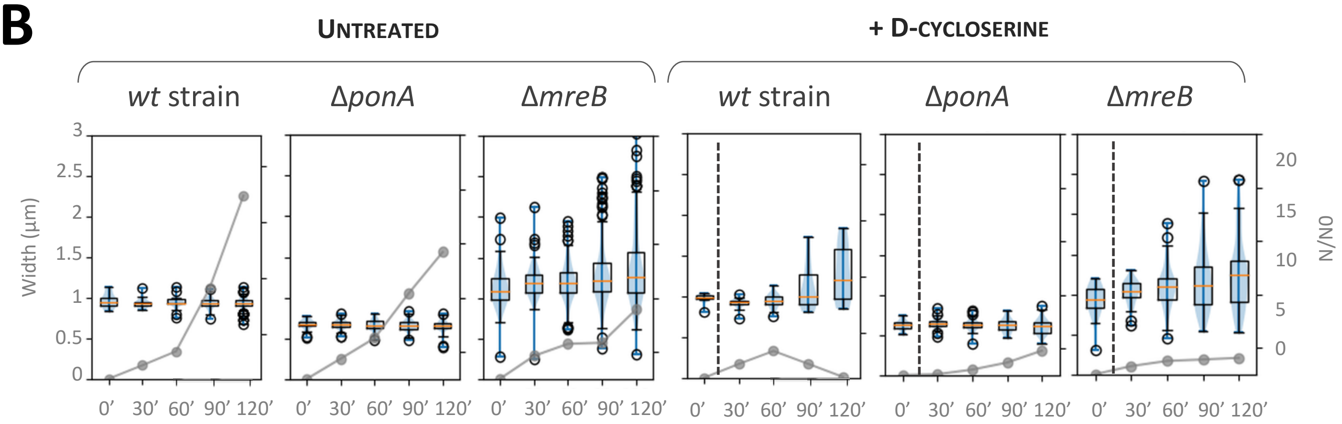


**Fig. S10 - Growth and bulging of D-cycloserine-treated cells depends on PBP1, encoded by *ponA***

**A.** Timelapses realized on the wt strain, *ΔponA* and *ΔmreB* mutants untreated and treated with 100 µg/mL D-cycloserine (scale bars: 5 µm). Cell were grown until exponential phase before being spotted on agarose pad prepared in LB (t0), antibiotic was added at 20 min. Acquisitions were taken in phase contrast and epifluorescence (membrane staining) microscopy every 2 min, 1 image every 60 min is presented here (scale bar: 5µm). See corresponding **Suppl. movies S14-S17**.

**B.** Quantification of the number of cells and of cell diameter were carried out on time-lapses microscopy movies. The evolution of the number of cells (N), normalized by the number of cells at t0 (N0) is shown as grey curves, corresponding to the right axis. These curves are a mean of 3 independent experiments. The left axis correspond to cell width (in µm), depicted as boxplots on the graphs. Antibiotic were added at 20 min (dashed line).

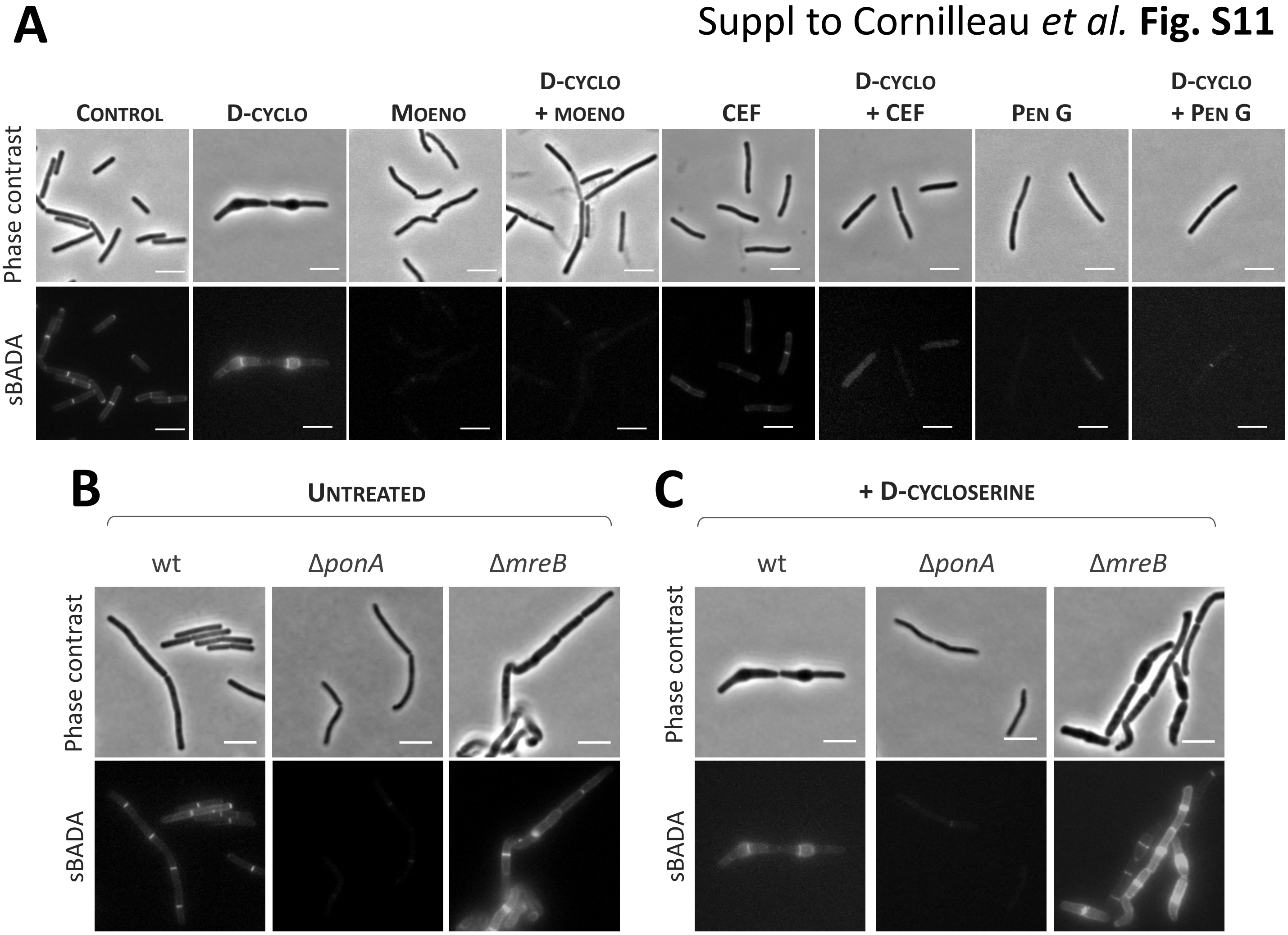


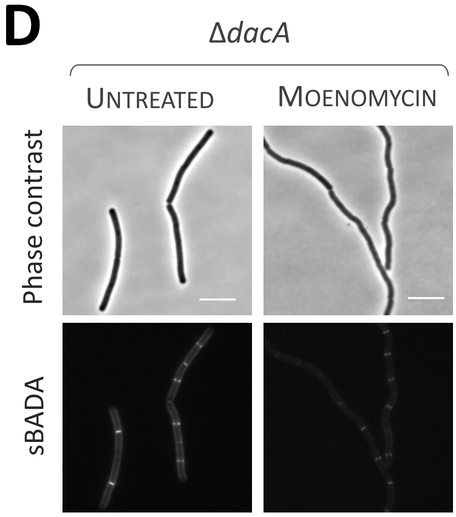


**Fig. S11 – Growth and bulging of D-cycloserine-treated cells depend on PBP1 activity**

Antibiotics were used at the following concentrations: 100 µg/mL D-cycloserine, 50 µg/mL moenomycin, 20 µg/mL penicillin G, 0.05 µg/mL cefuroxime.

**A.** Phase contrast and sBADA labelling of the wt strain. Cell were grown until exponential phase before addition of CW antibiotics and further incubation during 1 h. Same settings for all fluorescent images (scale bar: 5 µm).

**B.** Phase contrast and sBADA labelling of the wt strain, *ΔmreB, ΔponA* mutants. Same settings for all fluorescent images (scale bar: 5 µm).

**C.** Phase contrast and sBADA labelling of the wt strain, *ΔmreB, ΔponA* mutants treated with D-cycloserine. Cell were grown until exponential phase before addition of CW antibiotics and further incubation during 1 h. Same settings for all fluorescent images (scale bar: 5 µm).

**D.** Phase contrast and sBADA labelling of the *ΔdacA* mutant untreated and treated with moenomycin. Cell were grown until exponential phase before addition of CW antibiotics and further incubation during 1 h. Same settings for all fluorescent images (scale bar: 5 µm).
